# MUFASA: A Continuous-Time Stochastic Framework for Realistic Fluorescence Microscopy Simulation

**DOI:** 10.1101/2025.07.10.664068

**Authors:** Wessim Omezzine, Sébastien Schaub, Laure Blanc-Féraud, Luca Calatroni

## Abstract

We present MUFASA (**M**ulti-Protocol **U**nified **F**luorescence-based **A**dvanced **S**imulation **A**lgorithm), a physically grounded, continuous-time simulator for super-resolution fluorescence microscopy. By modeling fluorophore dynamics using continuous-time Markov chains, MUFASA’s simulation features yield realistic photon emission behavior across both Single Molecule Localization Microscopy (SMLM) and fluorescence fluctuation-based (FF-SRM) protocols—independently of frame duration and sampling. The framework supports both individual emitters and structure-level simulations, incorporating photophysical transitions, photobleaching, and camera properties.

To quantitatively validate simulations with real data, we introduce a novel validation metric based on the 1-Wasserstein distance between simulated and experimental photon-count distributions. In addition to simulation, another functionality estimates key photophysical parameters (e.g., molar extinction coefficient) and to suggest optimal light-source power ranges from fluctuation data. An intuitive Python-based graphical interface enables real-time parameter tuning, visualization, and TIFF export. Designed for biologists, physicists, microscopists, and numerical imaging engineers, MUFASA offers a practical platform for microscopy experiment design, hypothesis testing and the generation of realistic training data for data-driven microscopy methods across modalities.

## Introduction

Modern Super-resolution Microscopy (SRM) methods aim to overcome the diffraction limit by exploiting the photokinetics of the molecule and their intricate behaviors often represented in terms of electronic state transitions in Jablonski diagram (Figure 1).

**Figure 1:**
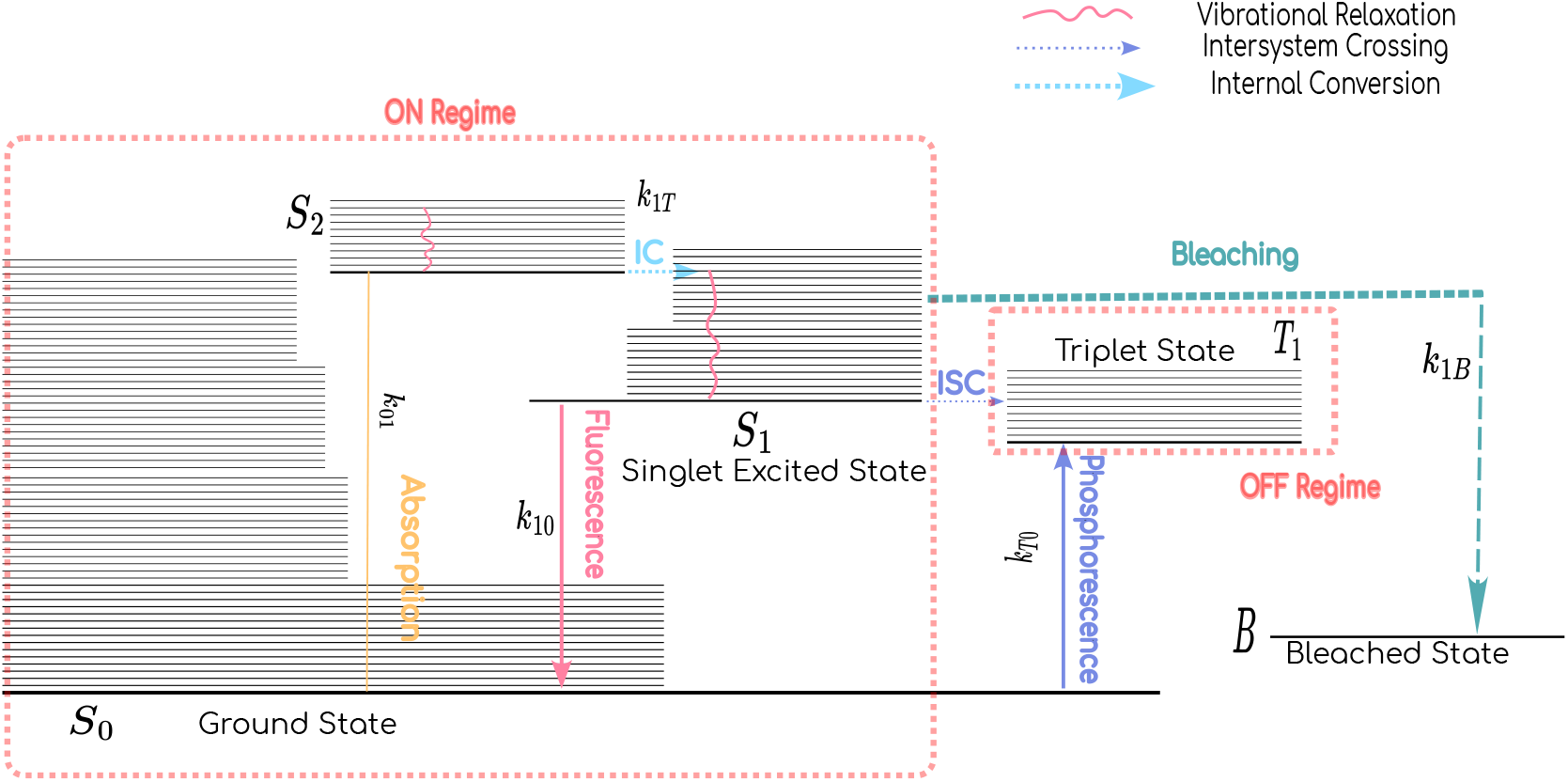
Fluorophore state transitions in fluorescence microscopy. The Jablonski diagram represents quantum states involved in photon absorption, emission, intersystem crossing, and photobleaching. Vertical solid arrows indicate radiative and non-radiative transitions between the ground singlet state (*S*_0_), the first excited singlet state (*S*_1_), and the triplet state (*T*_1_). Wavy and dotted lines represent non-radiative transitions. Additional transitions from a non-activated state (NA) and to a bleached state (B) are included to reflect specific behaviors observed in methods like PALM. This description of the state space forms the basis for the continuous-time Markov chain modeling used in MUFASA.

As outlined in [1], SRM can be grouped into three main families. Early advancements were made with STimulated-Emission-Depletion (STED) [2], which improves resolution by selectively deactivating fluorophores at the periphery of the excitation spot using a depletion laser, thereby reducing the effective point spread function. Structured Illumination Microscopy (SIM) [3] enhances resolution by projecting known light patterns and computationally reconstructing high-frequency information beyond the diffraction limit. Subsequently, Single Molecule Localization Microscopy (SMLM) [4] emerged as a prominent family of methods, enabling superresolution imaging through the precise localization of photoactivatable *blinking* fluorophores that stochastically switch between fluorescent (on) and nonfluorescent (off) regimes—a behavior commonly referred to as *fluorescence blinking*. By temporally isolating the emission of individual molecules, SMLM approaches such as STochastic Optical Reconstruction Microscopy (StORM) [5] and PhotoActivated Localization Microscopy (PALM) [6] can localize each emitter and reconstruct high-resolution images from these localizations. These methods typically operate under high light power conditions (up to several kW cm^−2^) to drive fluorophores into longlived dark states or induce controlled photobleaching, thereby temporally separating emitters.

Building on this perspective, Fluorescence Fluctuation-based SRM (FF-SRM) techniques [7] have been developed, which exploit the temporal intensity fluctuations of fluorophores to achieve resolution enhancement. Unlike SMLM, which isolates single emitters in time, FF-SRM analyzes the collective temporal behavior of multiple emitters within each diffraction-limited pixel. These methods usually operate under low laser power conditions (e.g., 0.1 W cm^−2^), which preserve fluorophore activity and lead to mean continuous photon emission. These fluctuations are often modeled using Poisson statistics, under the assumption that photon emission is a memoryless process occurring with constant rate over short intervals [8]. Higher-order statistical moments (e.g., cumulants of order *N* ≥ 2) are then computed to suppress noise and enhance spatial detail. By leveraging these temporal dependencies, FF-SRM reconstructs super-resolved images without requiring explicit localization of individual molecules.

However, SRM still faces challenges in both reproducibility and interpretability. Imaging results are highly sensitive to experimental parameters and the fluorophores photophysical properties, which are often poorly characterized. For instance, small changes in laser intensity can drastically alter blinking behavior, leading to inconsistent reconstructions [9]. Misinterpretation of photophysical artifacts—such as fluorophore reactivation or triplet state transitions—can result in false-positive localizations or incorrect molecule counts [10, 11]. Physically grounded simulation frameworks would be an excellent tool not only to guide experimental design, enable protocol comparison, support method validation, and generate synthetic training data for datadriven methods. Physically grounded simulators can produce realistic fluorescence signals with known ground truth, estimating key photophysical effects such as blinking kinetics, bleaching, and noise. This enables training models that generalize better across imaging conditions while being benchmarked rigorously using synthetic validation datasets.

Multiple efforts have been made to simulate photophysics. Early tools such as SuReSim [12] and FluoSim [13] relied on simplified photoswitching models, primarily focusing on “on” and “off” regimes to simulate StORM and PALM dynamics, implying a constant photon flux during “on” regimes. The switching behavior is governed by *transition rates*– that is, the probabilities per unit time of transitioning between states. In these simulators, these transition rates were user-defined and not derived from first-principles or measured experimental parameters, limiting their physical interpretability. More recent tools like VirtualSMLM [14] and SMIS [15] have better physical consistency by linking transition rates to experimental parameters. However, these simulators use fixed discrete time grids, which are intrinsically unable to describe rapid transitions, i.e occurring at sub-frame timescales.

The SOFI Simulation tool [16] addresses this limitation for fluctuation-based models by employing Continuous-Time Markov Chains (CTMCs, rather than modeling fluorophore dynamics as a sequence of updates on a fixed temporal grid. This enables a more faithful representation of blinking dynamics, which occur randomly but with well-defined rates, independent of any external clock.

However, even this approach remains limited in scope. The SOFI Simulation tool applies the CTMC only to the switching between observable “on” and “off” regimes. Within the “on” regime itself, photon emission is still modeled as a constant-rate process, thereby ignoring the rapid quantum jumps between excited *S*_1_ and ground *S*_0_ singlet states that govern the actual photon emission events. Capturing these sub-frame dynamics requires extending the CTMC framework to model the full set of quantum state transitions involved in fluorescence.

To address this gap and encompass different fluorescence dynamics, we introduce the **M**ultiProtocol **U**nified **F**luorescence-based **A**dvanced **S**imulation **A**rchitecture **(MUFASA)**, a comprehensive, physics-inspired simulator that integrates both SMLM and FF-SRM protocols within a single continuous-time framework. Unlike previous tools that model either discrete switching events or simplified emission processes, MUFASA simulates quantum state transitions directly using CTMCs. This enables the accurate reproduction of both intermittent and continuous emission behaviors. MUFASA comprises two core features:

1. **MUFASA-Sim**, which performs protocol-aware simulations of photon emission based on quantum photophysics, including realistic transition times and photobleaching effects.
2. **MUFASA-Design**, which extracts relevant photophysical parameters–such as the molar extinction coefficient—from experimental fluctuation data, and provides optimal illumination power range for imaging.

MUFASA incorporates the detailed photophysical dynamics described by the Jablonski diagram, intp a comprehensive simulator, enabling the modeling of rapid transitions between ground and excited states. In addition to its simulation capabilities, MUFASA includes a graphical user interface (GUI) for interactive exploration and a quantitative validation framework for assessing the realism of generated data. Specifically, we introduce the use of the 1-Wasserstein distance—a statistical metric that quantifies the difference between probability distributions based on the minimal transport cost needed to transform one distribution into another. This enables rigorous comparison between simulated photon emission statistics and experimental single-molecule traces.

Designed for both flexibility and realism, MUFASA serves as a valuable tool for a large community of researchers involved in the design of SRM experiments, from experimental sample preparation to numerical simulation, such as hypothesis testing, or generating physically grounded training data and realistic simulations for data-driven SRM methods.

## Results

An overview of the MUFASA simulation framework and its core components is illustrated in Figure 2, which provides visual context for the analyses presented in the following subsections.

**Figure 2:**
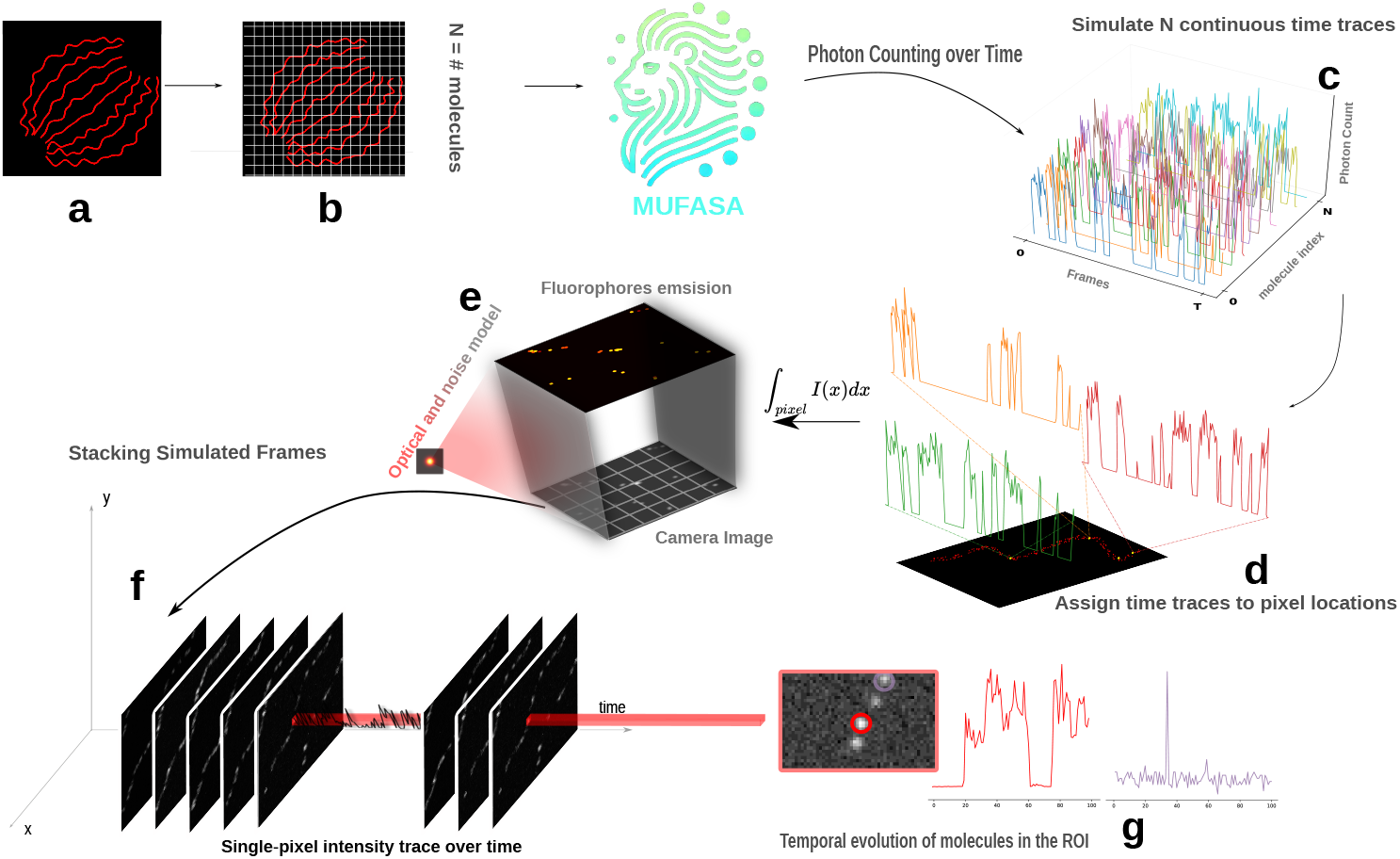
The MUFASA simulation pipeline for photophysics-aware fluorescence microscopy imaging. **(a)** A simulated biological structure is populated with discrete fluorophore labels to serve as the input to the MUFASA framework. **(b)** Fluorophore positions are discretized onto a regular 2D grid of *N* sampling sites. **(c)** Depending on the protocol, MUFASA generates *N* high-temporal-resolution photon-emission time traces by simulating its photophysical state transitions (excitation, emission, dark states) using a continuous-time Markov chain (CTMC) model and records the resulting per-molecule photon-count time traces. **(d)**Photon counts from all fluorophores within each grid cell are summed over all discrete time intervals to yield the total fluorescence contribution per pixel—i.e. the raw, per-frame emission signal. **(e)** Each raw frame is processed through a simulation of the microscope’s optical response: first, the high-resolution photon map is convolved with the point spread function (PSF) to model diffraction blur. Then, spatial undersampling is applied by summing photon counts within blocks of highresolution pixels to match the coarser detector grid. Finally, realistic camera noise is added (e.g., shot noise, EMCCD amplification), yielding a noisy, realistic resolution grayscale image stack. **(f)** The final output is a time-lapse series of low-resolution images that faithfully conveys both spatial heterogeneity and the temporal photophysics inherent to specific modalities. **(g)** A zoomed-in camera frame shows two selected fluorophores within a defined region of interest (ROI), marked by red and violet circles. For each fluorophore, the central pixel’s intensity is tracked over time under the simulated acquisition sequence, revealing temporal fluorescence traces (middle: red; right: violet). These traces reflect differences in photoswitching behavior and emission variability captured by MUFASA.

Under the SMLM-type dynamics, we simulate isolated fluorophores to examine protocol-specific emission statistics at high temporal resolution (Figure 2b). We qualitatively analyze the distribution of emitted photons over time, allowing us to study how varying experimental conditions influence molecular dynamics. These traces reveal protocol-dependent behaviors: continuous emission for FF, zero-inflated blinking in StORM, and sparse activation in PALM. We extended the framework to model the stochastic fluorescence dynamics of multiple independent fluorophores within a biological structure (Figure 2e). This extension builds on the single-molecule simulation pipeline (Figure 2a), but now requires additional input: the spatial coordinates of all fluorophores in the structure. Each fluorophore is then treated as an independent emitter, governed by the same photophysical parameters—such as state transition rates and excitation conditions—as in the single-molecule case. After simulating time dynamics, MUFASA models the optical system and the camera detection process (Figure 2c) using a point spread function (PSF) convolution followed by spatial downsampling and noise modeling. The PSF accounts for diffraction-limited optical blur, while the camera model incorporates detector-specific characteristics such as pixel size, shot noise and gain fluctuations, producing realistic low-resolution image frames. When applied to simulations of microtubule structures labeled with 6,498 fluorophores, MUFASA successfully reproduces contrast patterns and spatial density variations characteristic of each SRM modality, for instance closely mirroring observations from experimental SMLM acquisitions.

The MUFASA-Design feature complements MUFASA-Sim by estimating key photophysical parameters (e.g., molar extinction coefficients) from experimental data and simulating the optimal illumination power range, thereby streamlining experimental planning and optimization. Using enhanced green fluorescent protein (EGFP) and Alexa Fluor 647 as examples (see Supplementary Note 7), we show that the estimated parameters deviate by less than 10% from reference values.

To summarize, these results are validated across four SRM modalities that exploit temporal dynamics—FF, Blinking, StORM, and PALM—using both qualitative and quantitative benchmarks against experimental data. Further details are provided below.

### Simulating and approximating singlemolecule traces for different SRM protocols

Using MUFASA-Sim, we analyzed the continuous-time behavior of an isolated Alexa Fluor 647 molecule over 200 frames (10 ms per frame). For each frame, the number of emitted photons corresponds to the number of absorption–emission events, occuring within 1 − 2 nanoseconds as shown in Figure 3. This emission pattern directly results from the underlying quantum state dynamics illustrated in the same figure and formalized through transition rates in the CTMC model, as described in the Methods section on continuous-time dynamics.

**Figure 3:**
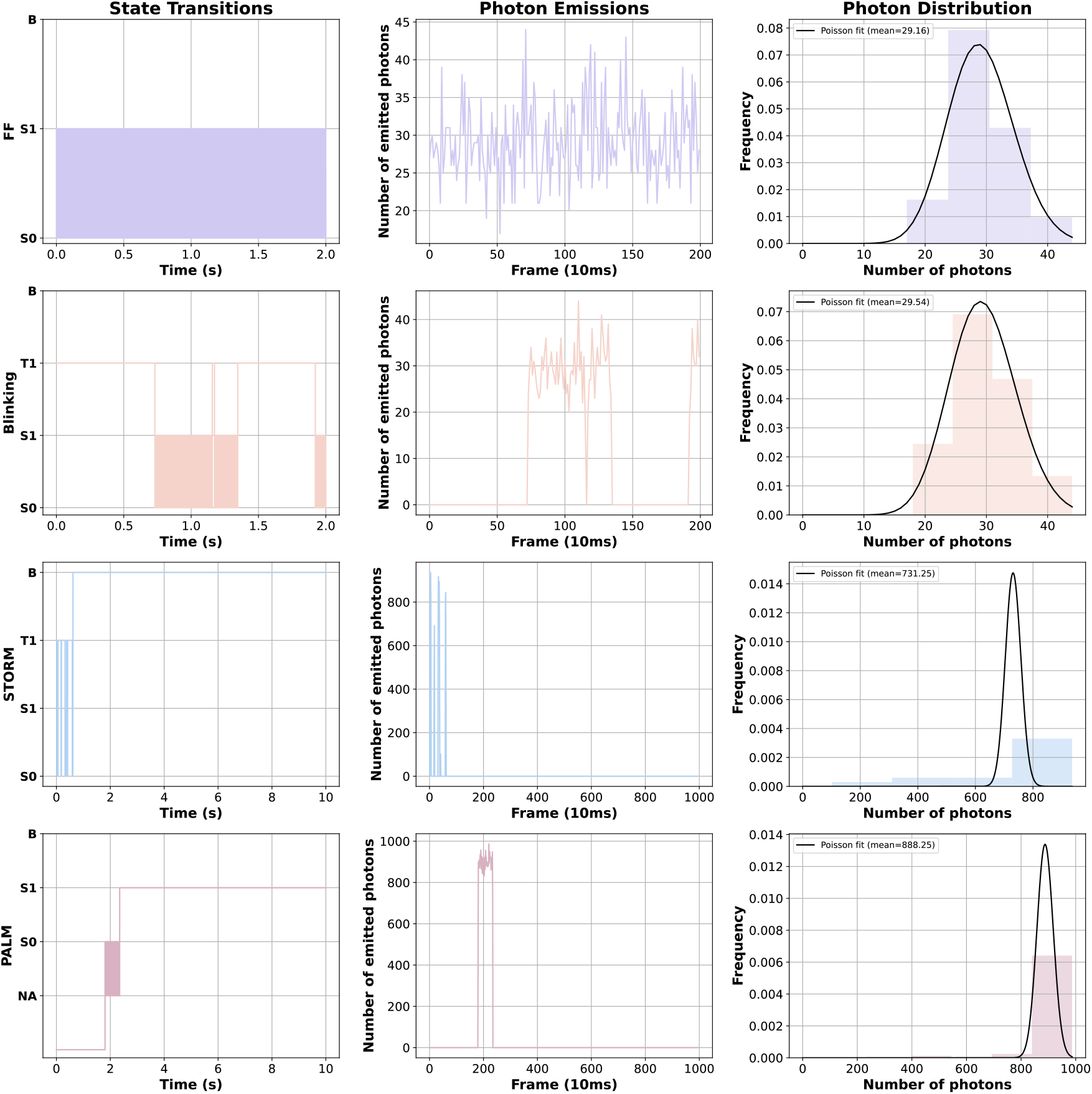
Comparative simulation of photophysical state transitions, single-molecule photon emission traces, and photon-count distributions across four fluorescence microscopy techniques: Fluorescence Fluctuation (FF), Blinking, Stochastic Optical Reconstruction Microscopy (StORM), and Photoactivated Localization Microscopy (PALM). The sampling frequency is fixed at 10 ms per frame for all protocols.

We simulated Alexa Fluor 647 fluorescence dynamics under four protocols: FF, Blinking (200 frames 10ms per frame), StORM and PALM (1000 frames, 10 ms per frame), using photophysical rates from the Thermo Fisher Scientific datasheet [17]. We included all states depicted in the Jablonski diagram (figure 1). Transitions between quantum states are governed by rate constants *k*_*ij*_, representing the probability per second of transitioning from state *i* to state *j*. To simplify the model without sacrificing accuracy, transitions with rates *k*_*ij*_ *<* 10^−4^*s*^−1^ were pruned. Protocol differences derive solely from variations in experimental conditions like illumination power (0.1 vs 5*kW/cm*^2^), intersystem crossing (ISC) rates, which modulate the probability of transitioning from the excited singlet state to the long-lived triplet state.

MUFASA-Sim is built on continuous-time Markov chains rather than discrete updates; hence, it faithfully captures rapid sub-frame transitions, where the lifetime of each state is randomly drawn from an exponential distribution, as shown in Figure 4. These transitions would not be captured by a frame-based update scheme.

**Figure 4:**
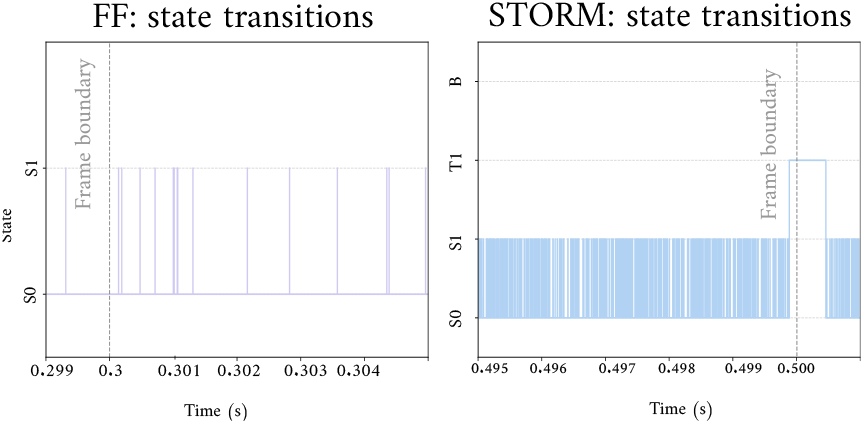
Sub-frame photophysical transitions revealed by MUFASA’s continuous-time CTMC model. Zoom in on single-molecule state trajectories under FF (left) and StORM (right) conditions, overlaid with the 10 ms frame boundaries (dashed line). MUFASA generates exponential holding time for states capturing rapid transitions, independently of frame duration.

Under the FF protocol, where the light power is relatively low (0.1 W/cm^2^), the molecule exhibits continuous emission, rapidly alternating between the ground and excited singlet states. For Alexa Fluor 647, the mean lifetime in the excited state (*τ*_1_ = 1 ns) is much shorter than the typical holding time in the ground state, which can reach up to 1 ms. The per-frame photon emission trace fluctuates around a mean of approximately 300 photons. A statistical analysis of the distribution of emitted photon counts reveals that it can be well-approximated by a Poisson distribution. Theoretical results derived in Section 2 show that, under certain conditions, the emission process indeed follows a Poisson distribution. More precisely, using Equation 1, one can compute *k*_01_ and find that the theoretical Poisson rate *µ*_theoretical_ matches the observed mean photon count.

The blinking and StORM protocol share similar photophysical mechanisms: both increase the probability of intersystem crossing (ISC) and extend the time spent in the triplet state. However, they differ significantly in terms of excitation power and the likelihood of photobleaching. StORM typically uses a higher laser intensity (5 kW/cm^2^), leading to a greater number of emitted photons during each activation event, compared to blinking (1 kW/cm^2^). In both cases, the molecule alternates between dark and fluorescent regimes, producing emission traces with prolonged zero-photon intervals during the off state.

PALM, in contrast, relies on the random activation of molecules from a non-activated (NA) state. Activation from this NA state occurs with a probability *p*_act_ per second. Once activated, Alexa Fluor 647 undergoes multiple absorption–emission cycles, typically within one or two frames (∼ 15 ms) before bleaching. Over 1000 frames (10 ms per frame), the zero-photon count fraction is 99.2%, with a heavy tail consisting of rare 1–2 frame bursts of high photon emission. In large experimental datasets, this pattern persists over tens or even hundreds of thousands of frames, with fluorescent spots generally lasting no more than 1–3 frames.

Across Blinking, StORM, and PALM protocols, the emission traces reveal features such as activation events, long triplet-state dwell times, and sparse burst-like behavior. These characteristics lead to deviations from the standard Poisson assumption, often producing zero-inflated photon count distributions. More appropriate models, such as Poisson–Gamma (negative binomial) mixtures, may better capture the increased variance introduced by blinking and state switching. However, rather than relying on protocol-specific statistical models, MUFASA addresses this variability through a unified framework based on continuous-time Markov chains (CTMCs), enabling realistic simulation of photophysical state transitions across all SRM protocols.

### MUFASA-Sim: Biological structure simulation

We extend the use of MUFASA-Sim beyond singlemolecule modeling to biological structures for a given spatial pattern.

As a test, we used a synthetic microtubule derived from fluorophore coordinate data available in [18] to generate protocol-specific image sequences. The geometry, illustrated in Supplementary Figure 4, comprises a dense distribution of 6,498 fluorophores arranged over a 2048 *×* 2048 grid. Local molecular densities vary between 0 and 4 emitters per pixel, mimicking experimentally observed heterogeneity. To capture realistic temporal behavior, highresolution photon emission traces were generated by simulating the continuous-time dynamics of fluorophores using protocol-specific CTMC models. These emission maps were then blurred by a simplified Gaussian point spread function (PSF)—noting that more advanced PSF models can be incorporated, as described in [19], downsampled by a factor *L* = 4, and passed through a noise-aware camera model to produce digitally quantized output. The noise model includes Poisson-distributed shot noise, additive Gaussian readout noise, and EM gain fluctuations according to models established in [20–22]. Acquisition parameters, including optical and camera properties, are summarized in Supplementary Table 1, with protocol-specific experimental and molecular details provided in Supplementary Tables 2 and 3, respectively.

**Table 1:**
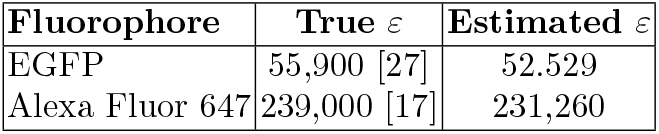
Comparison of true and MUFASA-Design estimated molar extinction coefficients.

| Fluorophore | True $\varepsilon$ | Estimated $\varepsilon$ |
| --- | --- | --- |
| EGFP | 55,900 [27] | 52.529 |
| Alexa Fluor 647 | 239,000 [17] | 231,260 |

Protocol-specific simulations are shown in Figure 5 and Supplementary Figures 5–9, each demonstrating distinct behavior. All simulations used the same fluorophore model; environmental parameters such as excitation intensity and non-radiative decay rates were adjusted to reflect the operational principles of each SRM modality. The parameters used for the camera models are detailed in Supplementary Note 5.

**Figure 5:**
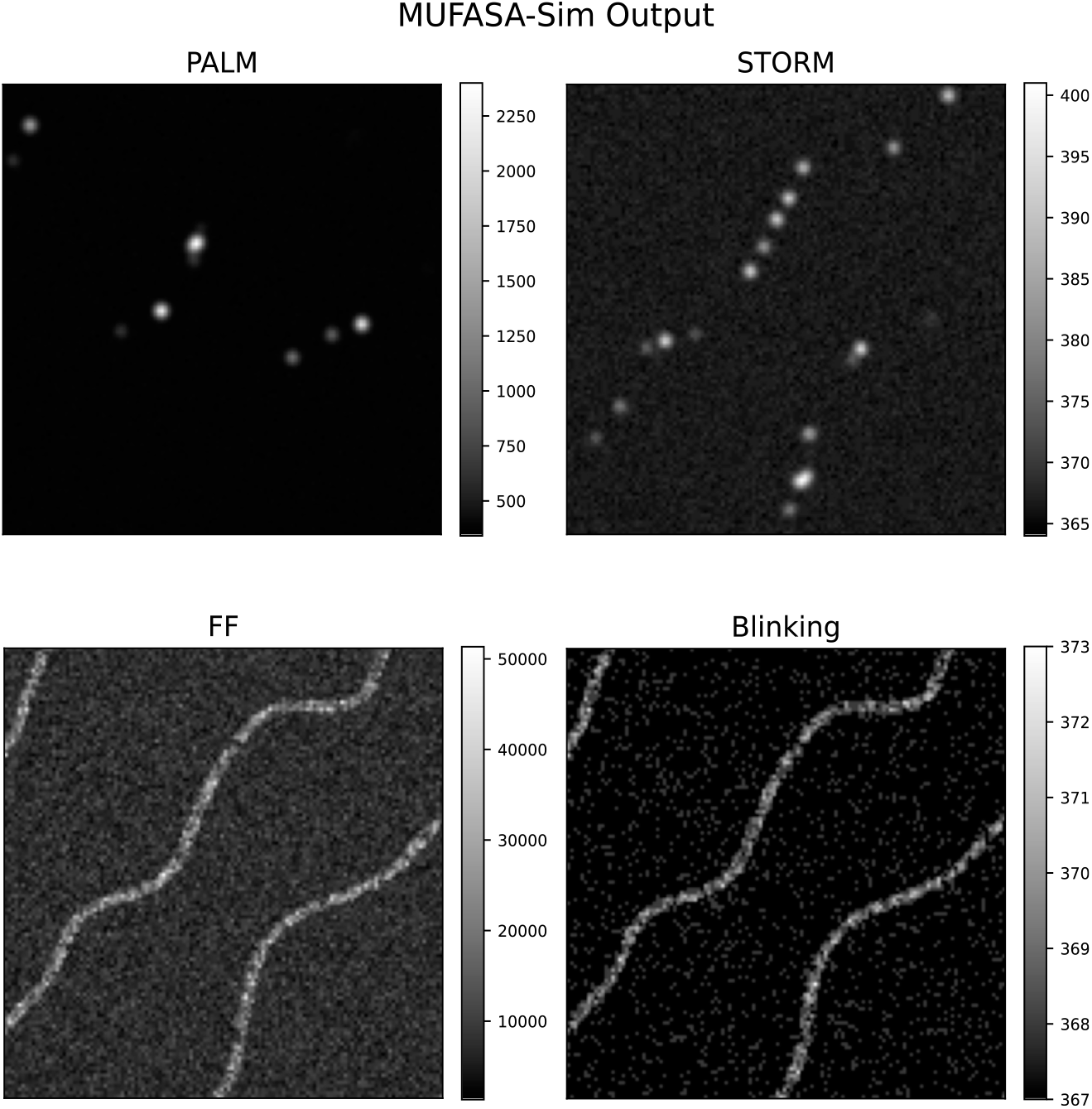
Protocol-specific simulated acquisitions of microtubule structures using MUFASA-Sim. Representative output frames from simulations of a microtubule bundle under four distinct SRM protocols: PALM, StORM, FF, and Blinking. All pixel intensities are expressed in analog-to-digital units (ADU), as output by the simulated camera. Although the same fluorophore model is used across all conditions, protocol-specific behaviors are recapitulated through changes in excitation regime, transition dynamics, and detection noise. PALM and StORM produce sparse, high-intensity single-molecule events due to activation and rapid photobleaching. FF imaging yields dense, continuous emission with spatially structured signal, while Blinking exhibits temporally modulated sparsity reflecting stochastic transitions between emissive and dark states.

These qualitative visual results confirm that MUFASA-Sim faithfully reproduces the protocol-specific behaviors of different SRM techniques. Quantitative validation against experimental data is presented in the Validation Section.

### MUFASA-Design: Estimation of molar extinction coefficient *ε*

The molar extinction coefficient *ε* quantifies how efficiently a molecule absorbs light at a given wavelength. This parameter is important for biologists, as it accurately quantifies concentration [23]. For physicists, it plays a central role in modeling light–matter interactions in nanophotonics and quantum optics [24–26].

However, determining *ε* is challenging because it is influenced by the fluorophore quantum yield—the probability that an absorbed photon results in the emission of a new phton—which itself depends on environmental factors such as solvent polarity, pH, and temperature. Variations in quantum yield can lead to significant deviations in the observed absorbance, complicating accurate estimation of *ε* from experimental data [23].

MUFASA-Design enables the estimation of *ε* from fluorescence fluctuation data by inverting a physically grounded expression for the excitation rate *k*_01_ given by the Equation 1:

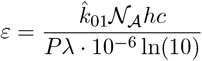

Here, 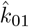 is the excitation rate estimated from fluorescence time traces after denoising the experimental signal (as detailed in Supplementary Note 3), 𝒩_*A*_ is Avogadro’s number, *h* is Planck’s constant, *c* is the speed of light, *P* is the excitation power density, and *λ* is the excitation wavelength.

We validated this approach by estimating its value for two widely used fluorophores (Table 1). In both cases, MUFASA-Design yielded estimates within 10% of the literature values, despite operating on noisy simulated emission data.

### MUFASA-Design: Illumination power range calibration

Proper calibration of laser light is a critical step in fluorescence microscopy, particularly in superresolution applications where photobleaching can severely limit data quality and experiment duration. In practice, the selection of illumination intensity often relies on trial and error, leading to potential overexposure or inadequate signal [28].

To address this, MUFASA-Design enables users to simulate the photokinetics of fluorophores across a wide range of light intensities. We applied the simulator to model EGFP behavior for excitation intensities ranging from 0.1 to 10 kW/cm^2^ under the FF protocol. For each setting, MUFASA-Design computes the expected photon emission per molecule.

These simulations offer a practical tool for calibrating illumination intensity in SRM acquisitions by enabling users to predict fluorophore response prior to experimentation. For example, MUFASA-Design simulations of EGFP photodynamics (Figure 6) reveal that at intermediate light powers (e.g., between 0.5 and 1 kW/cm^2^), one can achieve stable emission exceeding 3000 photons per frame for over 500 frames before photobleaching becomes relevant. This allows experimentalists to select illumination settings that balance brightness and longevity, reducing photodamage and avoiding suboptimal conditions. Rather than relying purely on experimental trial and error, MUFASA thus provides a principled approach to explore and constrain viable illumination power regimes before real sample acquisition.

**Figure 6:**
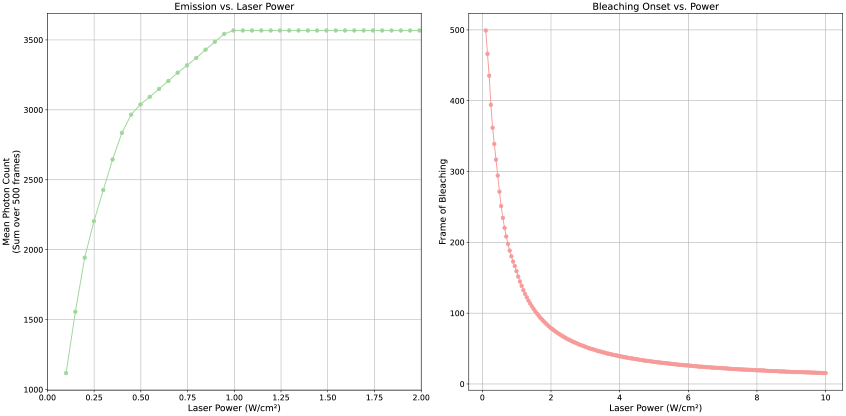
Simulation of EGFP photodynamics across illumination powers using MUFASA-Design. **Left:** Mean cumulative photons that a single EGFP molecule can emit before reaching the bleaching criterion when illuminated continuously with 488-nm light between 0.1 ™ 2kW/cm^2^. The initial linear rise ≤ 0.3kW/cm^2^ reflects the excitation-limited regime, whereas the shoulder above (≈ 0.7kW/cm^2^) marks the cycle-limited behavior. **Right:** Expected frame index at which the bleaching event occurs for the same simulations. Because bleaching is modelled as a fixed maximum number of excitation cycles, increasing light power shortens the time required to reach this limit. This curve allows users to identify illumination regimes that ensure stable emission while minimizing photobleaching.

### Validation

In this section, we report the numerical assessments performed on MUFASA’s ability to replicate the stochastic dynamics of fluorophore emission by comparing it with experimental photon emission traces. As a benchmark, we consider SOFI-Simulation Tools [16], a widely used simulator based on fluorescence fluctuation analysis. This comparison focuses on two key imaging regimes: Blinking and Fluorescence Fluctuation (FF) microscopy. To quantitatively assess the effectiveness of the simulator in generating data with distributions similar to the observed ones, we used first-order Wasserstein distance [29], which captures differences in the overall shape and structure of the photon count distributions between simulated and experimental traces as detailed in Supplementary Note 9.

#### Comparative analysis and fluctuation-based statistical evaluation

SOFI Tools assumes a fixed emission rate during each “on” period, resulting in uniform photon plateaus across molecules. This tool is detailed in Supplementary Note 4. In contrast, MUFASA simulates continuous-time quantum state transitions, allowing emission rates to vary dynamically within “on” regimes.

The figures 7a and 7b display the differences in photon emissions simulated for three molecules. We analyze how the total photon count evolves as the molecules bleach over time. For each simulator, we generated three independent photon time traces over 3,000 frames. The first fluorophore was configured to bleach around frame 1,000, the second around frame 2,000, and the third remained stable throughout the entire acquisition. We then summed the emission traces to yield the total photon count over time, in order to highlight how each model handles photon emission dynamics during the “on” regime. The shaded regions represent the amplitude of emission fluctuations within each plateau.

**Figure 7:**
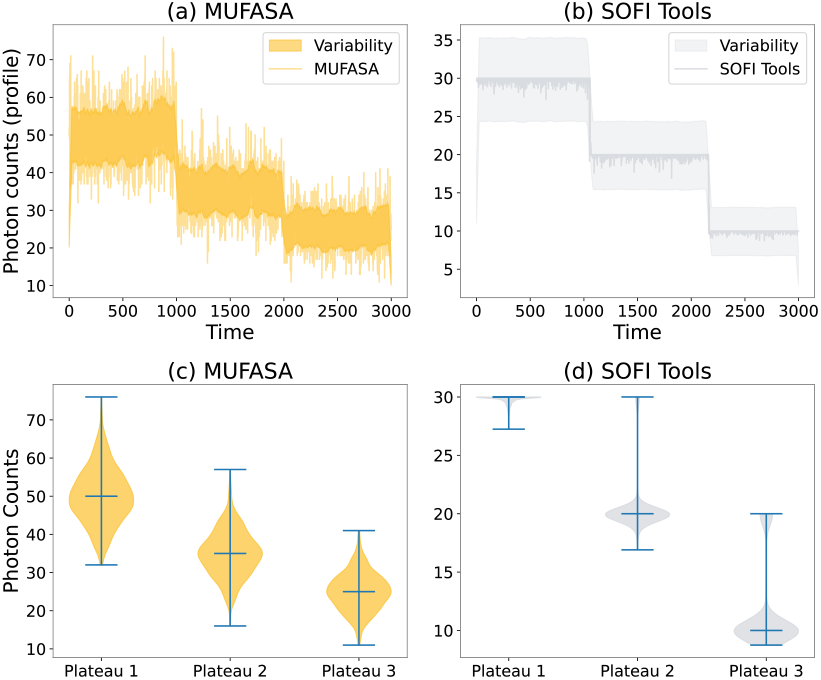
MUFASA captures sub-state variability in photon emission dynamics. **(a–b)** Photon emission traces from three fluorophores under the FF protocol, simulated with MUFASA (a) and SOFI Tools (b). Shaded regions indicate intra-plateau variability. MUFASA reveals greater fluctuations, suggesting more informative sub-state dynamics during the on-regime. **(c–d)** Violin plots of photon counts per emission plateau. MUFASA displays broader distributions, while SOFI Tools produces narrower, unimodal profiles due to its assumption of constant emission rates.

MUFASA exhibits greater variability compared to SOFI Tools, reflecting its explicit modeling of stochastic transitions between the ground state *S*_0_ and the singlet excited state *S*_1_. This variability is further illustrated in the violin plots (Figures 7c and 7d), which summarize the photon count distributions across the three plateaus.

MUFASA’s wider distributions capture the intrinsic randomness of fluorophore dynamics, in contrast to SOFI Tools, which assumes a constant emission rate and yields smoother traces with narrower distributions. This assumption neglects sub-state transitions within the “on” regime. MUFASA incorporates these transitions, allowing the detection of additional states and producing variable, non-uniform photon emission profiles even during the emitting regime.

### Quantitative assessment of StORM dynamics

To validate MUFASA under the StORM protocol, we compared its simulated photon emission traces to experimental data from [30]. For consistency, the analysis focused on the final 5000 frames of acquisition, where single-molecule events dominate due to widespread bleaching. We note that we validate MUFASA, under the FF protocol, against experimentally measured data in Supplementary Note 10.

We replicated acquisition conditions based on the metadata associated with the StORM dataset [30], including reported parameters such as excitation wavelength (641 nm), illumination power (∼1–2 kW/cm^2^), and camera specifications (Andor iXon EMCCD, 100 nm effective pixel size). Transition rates not explicitly reported—such as intersystem crossing *k*_1*T*_ and triplet decay *k*_*T* 0_—were empirically tuned to match the temporal dynamics observed in the real data. A detailed summary of the parameter inference and matching procedure is provided in Supplementary Note 8.

Beyond qualitative visual agreement illustrated in the left column of Figure 8, the right column of the same figure shows a qualitative comparison between experimental and simulated traces. MUFASA accurately captures the sparse, burst-like emission events characteristic of StORM imaging. The baseline signal and event amplitude are consistent with the experimental profile, demonstrating MUFASA ability to reproduce the intermittent, high-intensity emission typical of isolated fluorophores.

**Figure 8:**
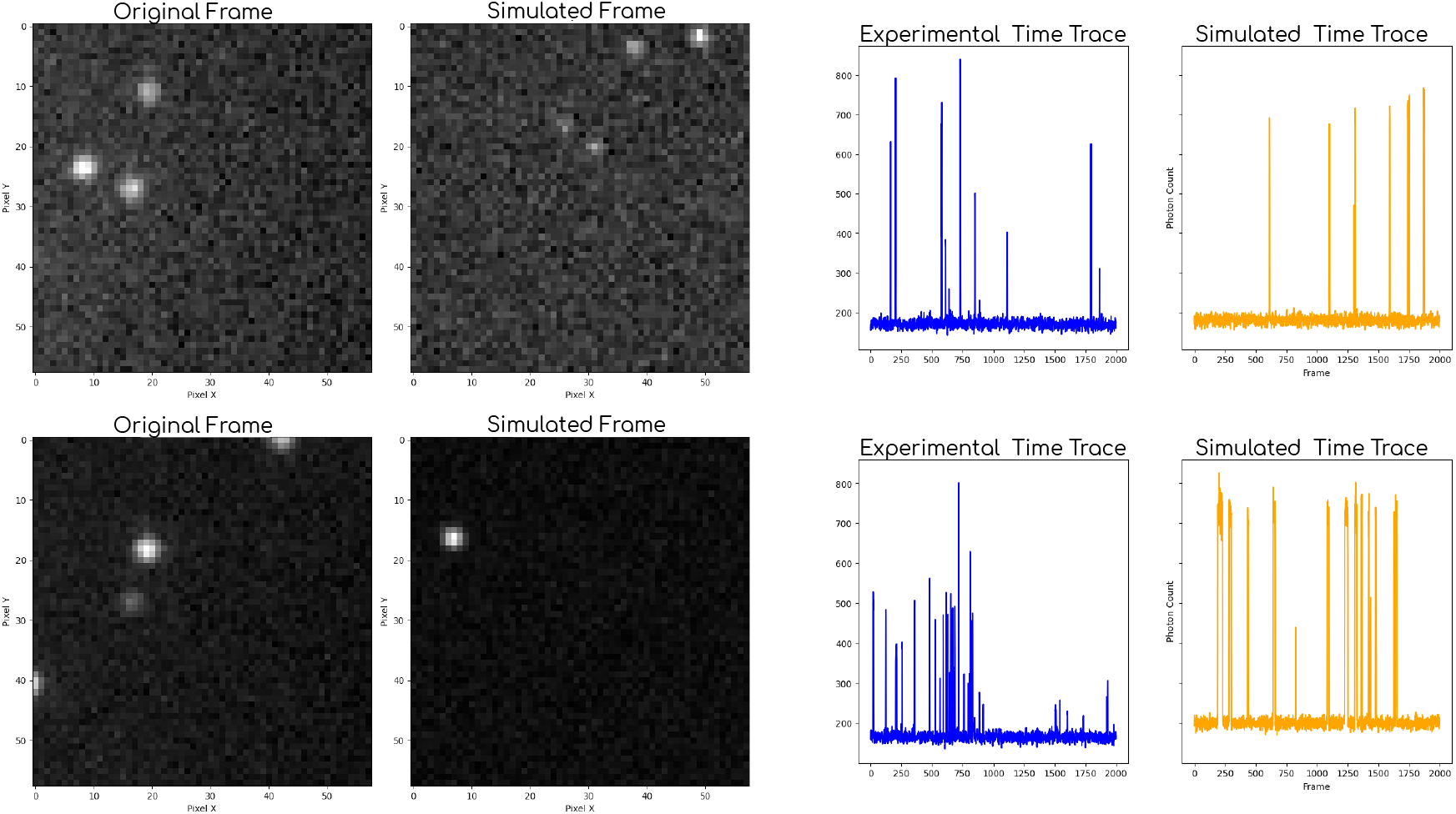
Validation of MUFASA under the StORM protocol: On the left: Each row shows a different time frame: left column corresponds to real experimental acquisitions, and right column displays the corresponding MUFASA-Sim outputs under matched conditions. **On the right:** Comparison of two photon count time traces between experimental StORM data (left, blue) and MUFASA simulations (right, orange). MUFASA reproduces the burst-like, high-intensity events and stable background observed in real single-molecule datasets. Traces shown correspond to the final 2000 frames of acquisition, where spatial and temporal sparsity dominates.

**Figure 9:**
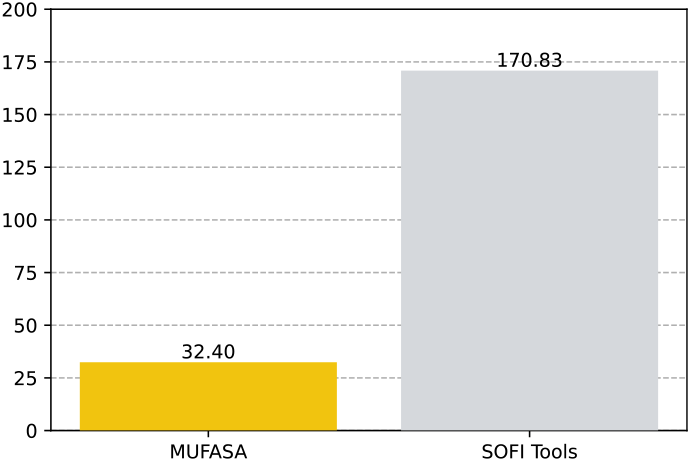
Comparison of simulation fidelity across models using the Wasserstein distance. Wasserstein distances (*W*_1_) between experimental photon count distributions and simulation models: MUFASA and SOFI Tools. Lower values indicate closer alignment with the empirical distribution. MUFASA achieves the best match, highlighting its capacity to capture realistic emission variability under the StORM protocol.

This alignment is confirmed with a Wasserstein distance (*W*_1_ = 32.40) between MUFASA simulations and the experimental data, significantly lower than SOFI Tools (*W*_1_ = 170.83).

## Discussion

MUFASA brings together simulation, quantitative validation, and experimental design in a unified framework for the description of fluorescence microscopy dynamics. At its core, it is a continuoustime Markov model that drives both single-molecule and structure-level simulations. This foundation not only enables realistic modeling of fluorophore dynamics (MUFASA-Sim), but also illumination powers MUFASA-Design—our dedicated module for estimating photophysical parameters such as the molar extinction coefficient *ε*.

This mathematical underpinning allows us to infer *ε* from experimental fluctuation data, a task traditionally hampered by environmental sensitivity and the absence of ground-truth rates. By estimating the excitation rate *k*_01_ from denoised traces, we sidestep the need for bulk measurements and offer a more direct, physically interpretable estimation pipeline.

MUFASA is also built for flexibility across protocols. It reproduces both sparse emission patterns typical of SMLM methods like StORM and PALM, and the high-density, continuous dynamics seen in FF-based techniques. This is made possible by a modular transition graph structure that adapts to each protocol photokinetics—rather than relying on hand-tuned switching assumptions.

A gap in previous simulation frameworks was the lack of rigorous validation: Virtual SMLM [14] relied primarily on visual matching of StORM images, without statistical comparison to experimental data, while SOFI Simulation Tools [16] focused on reconstruction quality using SOFI-specific metrics, without assessing the fidelity of raw photon emission dynamics. Here, we introduce the use of the 1-Wasserstein distance to quantitatively compare emission distributions from simulated and experimental datasets. This metric captures not only mean differences but also the geometric shape of distributions—offering a more appropriate evaluation metric than classical ones.

That said, we also acknowledge the limitations. While MUFASA better matches experimental traces than other simulators, it does not achieve perfect modeling. This discrepancy comes from several sources: camera effects in MUFASA are modeled using standard models, without full calibration to specific hardware; the point spread function (PSF) is over-simplified or borrowed from available datasets, introducing further sources of errors.

More importantly, MUFASA is not designed to replicate individual experiments. It does not simulate molecular diffusion or motion within the sample—fluorophore positions are assumed to be fixed and static. Nor does it capture the full environmental complexity that governs transition rates, such as pH, temperature, or oxygen concentration, which are often not available from experimental datasets. Instead, MUFASA is intended as a pre-experiment planning tool or a data generation engine. Its primary use cases include predicting how experimental parameter choices might impact imaging outcomes. Moreover, it can be used to and generate realistic synthetic data for training deep learning based models such as FluoGAN [31], DeepStORM [32] and DECODE [33] for image reconstruction.

Finally, the architecture is modular by design, allowing additional imaging modalities to be incorporated without modifying the core simulation logic. For instance, extension to three-dimensional (3D) structures is straightforward; because photon emissions are modeled independently of spatial arrangement, simulated traces can be assigned to fluorophores distributed in 3D. Similarly, multicolour imaging can be enabled by assigning distinct excitation and emission properties to different fluorophore types, each governed by its own dynamical parameters. As data-driven methods and largescale analysis become more prevalent, MUFASA can serve not only as a simulator, but also as a physically grounded prior for inverse modeling, uncertainty quantification, and hypothesis testing. It can be used to generate realistic synthetic datasets for training data-driven models or to constrain machine learning pipelines with physically plausible priors.

By uniting simulation fidelity, statistical validation, and experimental design support, MUFASA lays the groundwork for more interpretable, reproducible, and optimized imaging pipelines.

## Methods

### Photoswitching as continuous-time Markov chain

In fluorescence microscopy, the stochastic absorption and emission of photons cause observable fluorescence fluctuations. Inspired by prior approaches such as as Photo-switching hidden Markov model (PSHMM) [34], Single-Molecule Imaging Simulator (SMIS) [15] and SOFI Simulation Tools [16], we define the molecule’s state as a continuous-time stochastic process *X*(*t*), with *t* ∈ [0, *T*] representing time over the duration of the experiment *T* and *X*(*t*) assuming values from discrete set of quantum states; more details are given below.

Assuming the Markov property -that future states depend only on the current statewe model, in fact, *X*(*t*) as a time-homogeneous *Continuous-Time Markov Chain* (CTMC). In CTMCs, state transitions occur at any time point, with holding times in each state following an exponential distribution. A justification of this assumption can be found in Supplementary Note 1.

#### State spaces

In fluorescence microscopy, “On” and “Off” states are commonly used to describe fluorophore behavior. We propose to associate these notions to the notion of ‘regimes’ to better capture the detailed nature of the transitions. During the “On” regime, a molecule cycles between the ground state *S*_0_ and the excited singlet state *T*_1_. The emission of one photon corresponds to the transition from *S*_1_ to *S*_0_. Inspired by the Jablonski diagram 1, we then define the state space of the CTMC model and denote it as 𝒮. The state space varies depending on the specific superresolution protocol. The general state space is thus 𝒮 = *{NA, S*_0_, *S*_1_, *T*_1_, *B}*, where *NA* represents the non-activated state, *S*_0_ and *S*_1_ correspond respectively to the ground and excited singlet states, *T*_1_ denotes the triplet state, and *B* is the irreversible bleached state.

Models in SMIS [15] and PSHMM [34] introduce multiple dark states including triplet or redox states, resulting in *N* distinct dark states. However, as noted in Remark 1 in [35], in the context of Markov chains, states with identical transition rates can be combined into a single state without losing the Markov property. This allows us to aggregate various dark states into a single regime, termed *T*_1_, which is accessible exclusively from *S*_1_

Given state space 𝒮, we can associate it to a specific transition graph like in figure 10. These diagrams are inspired by prior formulations of fluorophore dynamics using hidden Markov models [34], which we generalize here to accommodate a range of microscopy protocols including StORM, PALM, and FF-based imaging. Note that, depending on the specific experimental conditions or photophysical model under consideration, only a subset of states in *S* may be relevant. This modularity enables the model to adapt flexibly to a wide range of protocols and fluorophore behaviors.

**Figure 10:**
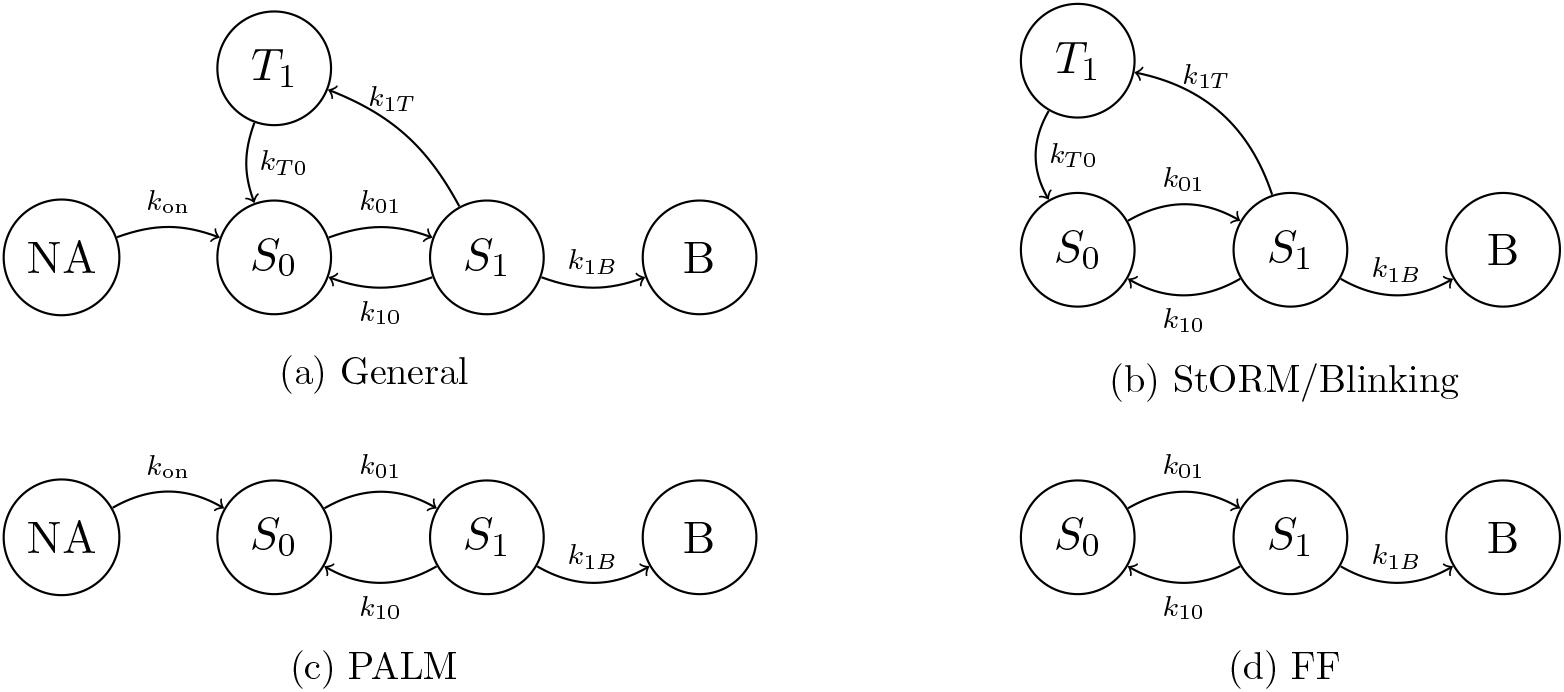
Quantum state transition diagrams for fluorophore modeling across microscopy protocols. Each diagram represents a continuous-time Markov model over discrete photophysical states, with arrows indicating allowed quantum transitions. (a) General five-state model incorporating all relevant transitions: activation from a non-active state (NA), cycling between ground *S*_0_ and excited *S*_1_ states, intersystem crossing to triplet *T*_1_, and irreversible bleaching (B). (b) Subset used for StORM and Blinking simulations. (c) PALM-specific model including activation from NA. (d) Model for fluctuation-based (FF) microscopy, assuming continuous excitation without activation or triplet transitions. All states are time-homogeneous and transitions follow exponential dwell-time assumptions.

#### Transition rates

The CTMC models we consider are defined in terms of environment-dependent transition rates which depend on specific properties of the considered molecules.

We mainly focus on the absorption/excitation rate *k*_01_ that governs the transition from *S*_0_ to *S*_1_. This rate can be linked to specific molecules and the laser properties through:

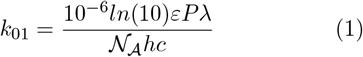

where *ε*[*M*^−1^*cm*^−1^] is the molar extinction coefficient of the molecule, *λ* [nm] is the excitation wavelength. Here, *P* [*W ·* −2] is the illumination power density at (*x, y*). 𝒩_*A*_ is the Avogadro number, *h* is Planck’s constant, and *c* is the speed of light [36]. This equation reveals that excitation becomes more probable for fluorophores with higher *ϵ* or stronger illumination *P*. It thus provides a direct and interpretable relationship between experimental parameters and the molecular transition dynamics that govern fluorescence.

Other transition rates in the CTMC model are expressed through the parameter *τ*, the mean holding times in given states, with 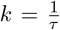. Specifically, the emission rate *k*_10_ from *S*_1_ to *S*_0_ is defined as 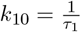 where *τ*_1_ ≈ 10^−9^*s* is the fluorescence lifetime. The intersystem crossing rate *k*_1*T*_ from *S*_1_ to *T*_1_, is instead given by 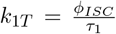, where the intersystem crossing quantum yield *ϕ*_*ISC*_ describes the sensitivity to molecular structures and environment. The triplet state decay is 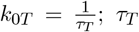 is significantly bigger than *τ*_1_ due to the spin-forbidden nature of the transition. The photobleaching rate *k*_1*B*_ representing the transition from *S*_1_ to the irreversible bleached state *B* is described by 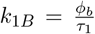, where *ϕ*_*b*_ represents the inverse of the average number of excitation-emission cycles before bleaching.

Defining precise closed-form expressions for these transition rates is challenging due to complex dependencies on environmental factors (temperature, pH, illumination intensity). Furthermore, the variability introduced by different experimental setups and environmental conditions prevents generalization across different scenarios. We provide general expressions that capture the fundamental behaviors without relying on overly specific parameters. For example, the bleaching rate *k*_1*B*_ is computed based on the expected number of excitation cycles a molecule undergoes before photobleaching. The intersystem crossing rate *k*_1*T*_ is set as a fixed fraction of the relaxation rate *k*_10_, reflecting the competition between radiative decay and triplet transition:

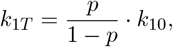

where *p* denotes the user-defined probability of triplet transition from the excited state. The return rate from the triplet state, *k*_*T* 0_, is defined from a user-specified triplet lifetime.

### Emission phenomena as a Poisson process

Fluorescence microscopy methods modeling photon emission typically assume a Poisson distribution for fluorescence fluctuations [35]. A fundamental justification for this hypothesis was presented by [37], where the authors showed that photon counts during the “on” regime can be modeled as a sum of Bernoulli trials—thus approximating a Poisson distribution in the limit where the number of emission attempts per interval becomes large, the probability of emission per attempt becomes small, and the expected number of emissions remains constant (i.e., *n* → ∞, *p* → 0, with *np* = *µ* fixed).

In this work, we provide a complementary justification for the Poissonian nature of fluorescence fluctuations using the continuous-time Markov chain (CTMC) framework described above. Our mathematical derivation is presented in full in Supplementary Note 2; it shows that under the realistic photophysical regime where excitation is much slower than relaxation, the photon emission CTMC process converges to a Poisson distribution without requiring the short-time approximation.

This connection between CTMC dynamics and Poisson statistics not only reinforces the modeling choices made in existing frameworks (see, e.g FluoGAN [31]), but also provides an interpretable physical mapping: : the reconstructed image intensity *x* corresponds to the mean photon count, *x* = *nk*_01_*t*, where *n* is the number of fluorescent molecules and *k*_01_ the excitation rate. This formulation embeds physical interpretability into the generative model, linking the learned image directly to the underlying photophysical parameters.

### Optical imaging system

The image formation process in fluorescence microscopy involves optical blur, spatial sampling, and the modeling of detector-specific noise. MUFASA simulates these effects through a modular pipeline, modeling both the optics and the camera electronics. In more details, emission at high spatial resolution is first convolved with a diffraction-limited point spread function (PSF), modeled for simplicity as a two-dimensional Gaussian. The resulting blurred image is then spatially undersampled by a factor *L >* 1, reflecting the realistic pixel size of the detector.

MUFASA-Sim further incorporates a detailed camera model, supporting three commonly used sensor architectures—CCD, EMCCD, and CMOS—based on established physical and statistical models from the literature [38, 39]. A complete description of the optical and electronic modeling pipeline is provided in Supplementary Note 3.

### Software architecture and performance

All simulations were run on a Linux workstation equipped with 16 physical CPU cores (22 logical), 30.8 GB of RAM, and Python 3.8. While GPU acceleration was not used, performance remained efficient thanks to multithreaded CPU execution and Intel MKL-optimized numerical libraries. Depending on the protocol and fluorophore density, simulation times ranged from less than 1 to 2 seconds per frame.

To quantify scalability, we benchmarked the runtime across protocols and increasing fluorophore counts. As shown in Supplementary Figure 9, simulation times grow approximately linearly with the number of molecules. StORM and PALM protocols exhibit lower runtime due to rapid photobleaching, whereas FF and Blinking simulations run longer due to persistent emission dynamics.

Importantly, MUFASA’s modular simulation architecture allows for efficient reuse of previously generated sequences. Simulated traces and emission profiles can be saved, recombined, or stacked across conditions—reducing redundant computations and enabling fast iteration during model testing or protocol design.

The graphical interface was developed using the PySide6 library [40]. The GUI is designed to support both reproducibility and hands-on exploration of simulation outputs. While optical effects like PSF blur are modeled using Gaussian kernels or userdefined profiles, MUFASA does not aim to replicate hardware-specific optics or cameras in fine detail. Instead, it provides a flexible, physically grounded platform for pre-experiment planning, data generation, and model testing.

## Code availability

The MUFASA simulation framework is openly available. The complete source code, including detailed documentation and a userfriendly graphical interface, is accessible at https://github.com/Joywessim/MUFASA_Fluorescence_Fluctuations_Simulation.

## Acknowledgements

This work is based upon work supported by the ANR JCJC project TASKABILE ANR-22-CE48-0010 and the funding received from the European Research Council (ERC) Starting project MALIN under the European Union’s Horizon Europe programme (grant agreement No. 101117133). Views and opinions expressed are however those of the author(s) only and do not necessarily reflect those of the European Union or the European Research Council Executive Agency. This work has been supported by the French government, through the 3IA Cote d’Azur Investments in the project managed by the National Research Agency (ANR) with the reference number ANR-23-IACL-0001.

Image acquisitions were conducted in the microscopy platform PIM (member of MICA microscopy platform and EMBRC-France), whose French state funds are managed by the ANR-10-INBS-0.

## References

[1] Alva, A. et al. Fluorescence fluctuation based super resolution microscopy, basic concepts for an easy start. bioRxiv (2022). URL https://www.biorxiv.org/content/early/2022/05/06/2022.05.06.490863. https://www.biorxiv.org/content/early/2022/05/06/2022.05.06.490863.full.pdf.

[2] Hell, S. W. & Wichmann, J. Breaking the diffraction resolution limit by stimulated emission: stimulated-emission-depletion fluorescence microscopy. Optics Letters 19, 780– 782 (1994).

[3] Bailey, B., Farkas, D. L., Taylor, D. L. & Lanni, F. Enhancement of axial resolution in fluorescence microscopy by standing-wave excitation. Nature 366, 44–48 (1993). URL 10.1038/366044a0.

[4] Jacquemet, G., Carisey, A. F., Hamidi, H., Henriques, R. & Leterrier, C. The cell biologist’s guide to super-resolution microscopy. Journal of Cell Science 133, jcs240713 (2020).

[5] Rust, M. J., Bates, M. & Zhuang, X. Subdiffraction-limit imaging by stochastic optical reconstruction microscopy (storm). Nature Methods 3, 793–795 (2006).

[6] Betzig, E. et al. Imaging intracellular fluorescent proteins at nanometer resolution. Science 313, 1642–1645 (2006).

[7] Dertinger, T., Colyer, R., Iyer, G., Weiss, S. & Enderlein, J. Fast, background-free, 3d super-resolution optical fluctuation imaging (sofi). Proceedings of the National Academy of Sciences 106, 22287–22292 (2009).

[8] Chen, Y., Müller, J. D., So, P. T. C. & Gratton, E. The photon counting histogram in fluorescence fluctuation spectroscopy. Biophysical Journal 77, 553–567 (1999). URL 10.1016/S0006-3495(99)76912-2.

[9] Nieuwenhuizen, R. P. et al. Quantitative localization microscopy: Effects of photophysics and labeling stoichiometry. PloS one 10, e0127989 (2015).

[10] Annibale, P., Vanni, S., Scarselli, M., Rothlisberger, U. & Radenovic, A. Quantitative photo activated localization microscopy: Unraveling the effects of photoblinking. PloS one 6, e22678 (2011).

[11] Lee, S.-H., Shin, J. Y., Lee, A. & Bustamante, C. Counting single photoactivatable fluorescent molecules by photoactivated localization microscopy (palm). Proceedings of the National Academy of Sciences 109, 17436–17441 (2012).

[12] Venkataramani, V., Herrmannsdörfer, F., Heilemann, M. & Kuner, T. SuReSim: simulating localization microscopy experiments from ground truth models. Nature Methods 13, 319– 321 (2016). URL 10.1038/nmeth.3795.

[13] Lagardére, M., Chamma, I., Bouilhol, E., Nikolski, M. & Thoumine, O. FluoSim: simulator of single molecule dynamics for fluorescence live-cell and super-resolution imaging of membrane proteins. Scientific Reports 10, 19954 (2020). URL 10.1038/s41598-020-77094-6.

[14] Griffié, J. et al. Virtual-smlm, a virtual environment for real-time interactive smlm acquisition. bioRxiv 2020.03.05.967893 (2020). URL https://www.biorxiv.org/content/early/2020/03/06/2020.03.05.967893. Preprint.

[15] Bourgeois, D. Single molecule imaging simulations with advanced fluorophore photophysics. Communications Biology 6, 53 (2023).

[16] Girsault, A. et al. SOFI Simulation Tool: A Software Package for Simulating and Testing Super-Resolution Optical Fluctuation Imaging. PLoS ONE 11, e0161602 (2016). URL https://journals.plos.org/plosone/article?id=10.1371/journal.pone.0161602. PMCID: PMC5008722.

[17] Scientific, T. F. The alexa fluor dye series (2024). URL https://www.thermofisher.com/fr/fr/home/references/molecular-probes-the-handbook/technical-notes-and-product-highlights/the-alexa-fluor-dye-series.html. Accessed: 16-Oct-2024.

[18] Sage, D. et al. Quantitative evaluation of software packages for single-molecule localization microscopy. Nature Methods 16, 387–395 (2019).

[19] Liu, Y. et al. Revisiting psf models: unifying framework and high-performance implementation (2025). URL https://arxiv.org/abs/2502.03170.2502.03170.

[20] Cox, G. & Sheppard, C. J. Shot noise, detection noise, and the limits of the light microscope. Journal of the Optical Society of America A 21, 820–827 (2004).

[21] Banterle, N., Bui, T.-H., Vojnovic, B., Collinson, L. M. & Kaminski, C. F. Camera-based image quality assessment for fluorescence microscopy. Journal of Microscopy 250, 35–49 (2013).

[22] Robbins, M. S. & Hadwen, B. J. The noise performance of electron multiplying charge-coupled devices. IEEE Transactions on Electron Devices 50, 1227–1232 (2003).

[23] Pace, C., Vajdos, F., Fee, L., Grimsley, G. & Gray, T. How to measure and predict the molar absorption coefficient of a protein. Protein Science 4, 2411–2423 (1995).

[24] Hertzog, M., Wang, M., Mony, J. & Börjesson, K. Strong light–matter interactions: a new direction within chemistry. Chemical Society Reviews 48, 937–961 (2019).

[25] Weight, B. M., Li, X. & Zhang, Y. Theory and modeling of light-matter interactions in chemistry: current and future. arXiv preprint 2303.10111 (2023).

[26] Kumar, A., Singh, S., Mudahar, G. S. & Thind, K. S. Molar extinction coefficients of some commonly used solvents. Radiation Physics and Chemistry 75, 731–736 (2006).

[27] Patterson, G. H., Knobel, S. M., Sharif, W. D., Kain, S. R. & Piston, D. W. Use of the green fluorescent protein and its mutants in quantitative fluorescence microscopy. Biophysical journal 73, 2782–2790 (1997).

[28] Boudreau, J., Paramasivam, G. & Godin, M. Excitation light dose engineering to reduce photobleaching and phototoxicity. Scientific Reports 6, 30892 (2016).

[29] Villani, C. Optimal transport: old and new, vol. 338 (Springer, 2008).

[30] Olivier, N. & Keller, D. STORM Vectashield datasets (Tubulin). 10.5281/zenodo.7620025 (2023). Zenodo dataset.

[31] Cachia, M., Stergiopoulou, V., Calatroni, L., Schaub, S. & Blanc-Féraud, L. Fluorescence image deconvolution microscopy via generative adversarial learning (fluogan). Inverse Problems 39, 054006 (2023). URL https://hal.science/hal-03790156v2.

[32] Nehme, E., Weiss, L. E., Michaeli, T. & Shechtman, Y. Deep-storm: super-resolution single-molecule microscopy by deep learning. Optica 5, 458–464 (2018).

[33] Speiser, A. et al. Deep learning enables fast and dense single-molecule localization with high accuracy. Nature Methods 18, 1082–1090 (2021).

[34] Patel, L. et al. A hidden markov model approach to characterizing the photo-switching behavior of fluorophores. Annals of Applied Statistics 13, 1397–1429 (2019).

[35] Staudt, T. et al. Statistical molecule counting in super-resolution fluorescence microscopy: Towards quantitative nanoscopy (2020). URL https://arxiv.org/abs/1903.11577.1903.11577.

[36] Avilov, S. et al. In cellulo evaluation of phototransformation quantum yields in fluorescent proteins used as markers for single-molecule localization microscopy. PloS One 9, e98362 (2014).

[37] Aspelmeier, T., Egner, A. & Munk, A. Modern statistical challenges in high-resolution fluorescence microscopy. Annual Review of Statistics and Its Application 2, 163–202 (2015).

[38] Tian, H. & El Gamal, A. Noise Analysis in CMOS Image Sensors. Dissertation, Stanford University, Stanford, CA (2000). Ph.D. dissertation.

[39] Hirsch, M., Wareham, R., Martin-Fernandez, M., Hobson, M. & Rolfe, D. A stochastic model for electron multiplication charge-coupled devices–from theory to practice. PLoS One 8, e53671 (2013). Epub 2013 Jan 31.

[40] Company, T. Q. Pyside6 - qt for python. https://doc.qt.io/qtforpython-6/(2024). xVersion 6.9.

