## Supplementary information for "MUFASA: A Continuous-Time Stochastic Framework for Realistic Fluorescence Microscopy Simulation"

This file contains Supplementary Notes, Figures, and Tables referenced in the main manuscript. We provide also two supplementary videos of two simulations.

#### Contents

**1 Supplementary Note 1: Continuous-time Markov Chains** **2**

**2 Supplementary Note 2: Approximation of CTMC with** $|\mathcal{S}| = 2$ **3**

**3 Supplementary Note 3: Optical and Camera System Modeling** **6**

**4 Supplementary Note 4: SOFI-Simlulation Tools** **9**

**5 Supplementary Note 5: Camera Configurations** **10**

**6 Supplementary Note 6: Experimental Protocol Parameters** **11**

**7 Supplementary Note 7: Extracting Single Molecule Signals from Experimental Acquisitions** **12**

**8 Supplementary Note 8: Experimental Conditions and Parameter Inference for STORM Validation** **14**

**9 Supplementary Note 9: Wasserstein Distance** $W_1$ **16**

**10 Supplementary Note 10: Quantitative assessments via Wasserstein-distance of FF dynamics** **17**

**11 Supplementary Figures** **19**

### 1 Supplementary Note 1: Continuous-time Markov Chains

In fluorescence microscopy, the stochastic nature of photon absorption and emission events leads to observable fluctuations in fluorescence signals. These fluctuations arise because photon-molecule interactions occur randomly over time, even under constant illumination. To model these dynamics, we represent the state of a fluorescent molecule as a stochastic process, denoted as $X(t) \in \mathcal{S}$, where $t$ represents continuous time and $\mathcal{S} \subset \mathbb{N} \cup \{0\}$ is the state space. Its general form is $\mathcal{S} = \{NA, S_0, S_1, T_1, B\}$, where $NA$ represents the non-activated state, $S_0$ and $S_1$ correspond respectively to the ground and excited singlet states, $T_1$ denotes the triplet state, and $B$ is the irreversible bleached state. Given that molecules have a finite set of quantum states, we can assume *i.e* $|S| < +\infty$.

We now introduce the equivalent in continuous time of the Markov assumption: the probability associated to the future state of the system within a certain time duration depends only on its current state, not on its past history, thus implying a memory-less property [9]. This applies for instance when the system interaction with its environment is minimal and the environment relaxes rapidly, losing its memory faster than the system dynamics, which is the case of molecular state transitions, which can often be effectively modeled as Markovian processes when all relevant states and interactions are accounted for [1].

Building upon the Markov property, we further assume that the process is time-homogeneous, meaning the transition probabilities between states are consistent over time. This leads us to model the fluorescence fluctuations using a Continuous-Time Markov Chain (CTMC) $(X(t))_{t\geq 0}$ with state space $\mathcal{S}$, [9]. By definition, the transition probabilities satisfy

$$\mathbb{P}(X(s+t) = j \mid X(s) = i) = \mathbb{P}(X(t) = j \mid X(0) = i) =: P_{ij}(t), \quad \forall s, t \geq 0, \ \forall i, j \in \mathcal{S}.$$

Unlike discrete-time Markov chains, where transitions occur at fixed time steps, CTMCs permit transitions at any continuous time point. The process remains in a given state $i \in \mathcal{S}$ for a random duration $T_i$, known as the *holding time*, which—through the memoryless property can be shown to follow an exponential distribution:

$$\mathbb{P}(T_i > t) = e^{-k_i t}, \quad t \geq 0,$$

where $k_i > 0$ denotes the total exit rate from state $i$.

The infinitesimal behavior of the CTMC is governed by the *generator matrix* (or rate matrix) $G = (k_{ij})$, whose entries satisfy:

$$k_{ij} \geq 0 \quad \text{for } i \neq j, \qquad k_{ii} = \mu_i = -\sum_{j \neq i} \mu_{ij}, \quad \forall\, i \in \mathcal{S}.$$

Thus, each row of $G$ sums to zero (differently from discrete-time Markov chains where transition matrices are stochastic), and the diagonal elements encode the total rate of leaving each state.

#### 2 Supplementary Note 2: Approximation of CTMC with $|\mathcal{S}| = 2$

The unified CTMC modeling used in MUFASA offers a theoretical insight into the interpretation of such parameters especially in the case of continuous emission before bleaching. We prove here that whenever the cardinality of the state space is equal to two and some assumptions on the transition rates hold, we can in fact approximate the Markov process by a Poisson process with a physical grounded parameter being its mean and variance.

Consider the simple example where $|\mathcal{S}| = 2$, with $\mathcal{S} = \{S_0, S_1\}$. The generator matrix of a CTMC process in this case reads:

$$G = \begin{bmatrix} -k_{01} & k_{01} \\ k_{10} & -k_{10} \end{bmatrix}$$

and the state transition matrix at time $t$ is given by definition by $P(t) = e^{Gt}$. We now assume

$$k_{01} \ll k_{10}.$$

This is a physically reasonable assumption in fluorescence photophysics, where excitation events are rare and relaxation from the excited state is typically fast [3].

For $t \in [0, +\infty]$, let $N(t)$ be the random process denoting the number of photons emitted in the interval $[0, t]$ and let us define $P(n, T) := \mathbb{P}\left(N(t) = n\right), \forall n \in \mathbb{N}^*$ be the probability of emitting exactly $n$ photons until instant $t$. Note that by definition there holds:

$$P(n, t) = \mathbb{P}(N(t) = n, X(t) = S_0) + \mathbb{P}(N(t) = n, X(t) = S_1),$$

which, using similar notations, can be decomposed as

$$P(n, t) = P_0(n, t) + P_1(n, t).$$

We study now the dynamics of the functions $P_i(n, t)$ in order to find its expression. For small time increments $\delta t$, we have:

$$\begin{aligned} P_0(n, t + \delta t) &= \mathbb{P}(N(t) = n, X(t + \delta t) = S_0) \\ &= \mathbb{P}(N(t) = n, X(t + \delta t) = S_0 | N(t) = n, X(t) = S_0)\mathbb{P}(N(t) = n, X(t) = S_0) \\ &+ \mathbb{P}(N(t) = n, X(t + \delta t) = S_0 | N(t) = n - 1, X(t) = S_1)\mathbb{P}(N(t) = n - 1, X(t) = S_1) \\ &= (1 - k_{01}\delta t)P_0(n, t) + k_{10}\delta t P_1(n - 1, t) + o(\delta t) \end{aligned}$$

We can thus compute the differential equation governing $P_0(n, t)$, which reads:

$$\frac{d}{dt}P_0(n, t) = -k_{01}P_0(n, t) + k_{10}P_1(n - 1, t)$$

Proceeding similarly for $P_1(n, t)$, we get the differential system:

$$\begin{cases} \frac{d}{dt}P_0(n, t) &= -k_{01}P_0(n, t) + k_{10}P_1(n - 1, t) \\ \frac{d}{dt}P_1(n, t) &= k_{01}P_0(n, t) - k_{10}P_1(n, t) \\ P_0(n, 0) &= \delta_{n0} \\ P_1(n, 0) &= 0 \end{cases} \tag{1}$$

where $\delta_{n0}$ denotes the Kronecker symbol. These equations are known as *master equations* [1]: equations for the probability that a stochastic particle can jump between different states is in one of these states at a time $t$.

To solve the master equations, we introduce the probability generating functions (PGFs):

$$\forall i = 0,1 \quad \forall z \ s.t \ |z| < 1, \ \ G_i(z,t) := \sum_{n=0}^{\infty} z^n P_i(n,t).$$

The PGF compactly encodes the entire desired probability distribution $P_i(n,t)$ into one single analytic expression. In more details, it transforms the infinite sequence of probabilities into a function whose behavior can be studied using differential equations. By multiplying both sides of the master equations (1) by $z^n$ and summing over all $n$, we get:

$$\begin{cases} \frac{d}{dt}G_0(z,t) & = -k_{01}G_0(z,t) + k_{10}zG_1(z,t) \\ \frac{d}{dt}G_1(z,t) & = k_{01}G_0(z,t) - k_{10}G_1(z,t) \\ G_0(z,0) & = \sum_{n=0}^{\infty} z^n P_0(n,0) = \sum_{n=0}^{\infty} z^n \delta_{n,0} = 1 \\ G_1(z,0) & = 0. \end{cases} \tag{2}$$

Our goal now is to solve this system for $G(z,t) = G_0(z,t) + G_1(z,t)$, from which we can compute the desired distribution. To do so, we now apply the Laplace transform to both PGFs to turn the system of time-domain differential equations into algebraic equations. The Laplace variable $s \in \mathbb{C}$ is assumed to lie in the domain where the integrals converge, which is typically $\mathrm{Re}(s) > 0$. We obtain:

$$\tilde{G}_i(z,s) = \mathcal{L}[G_i](z,s) = \int_0^{\infty} e^{-st} G_i(z,t)\, dt,$$

so that system (2) becomes:

$$\begin{cases} s\tilde{G}_0(z,s) - G_0(z,0) & = -k_{01}\tilde{G}_0(z,s) + k_{10}z\tilde{G}_1(z,s), \\ s\tilde{G}_1(z,s) - G_1(z,0) & = k_{01}\tilde{G}_0(z,s) - k_{10}\tilde{G}_1(z,s). \end{cases}$$

Now, since $G_0(z,0) = 1$ and $G_1(z,0) = 0$, after some algebraic manipulations we get:

$$\begin{cases} \tilde{G}_0(z,s) & = \frac{s+k_{10}}{s^2+s(k_{01}+k_{10})+k_{01}k_{10}(1-z)}, \\ \tilde{G}_1(z,s) & = \frac{k_{01}}{s^2+s(k_{01}+k_{10})+k_{01}k_{10}(1-z)}. \end{cases}$$

We thus deduce that:

$$\forall z \text{ s.t. } |z| < 1, \ \ \tilde{G}(z,s) = \tilde{G}_0(z,s) + \tilde{G}_1(z,s) = \frac{s + k_{10} + k_{01}}{s^2 + s(k_{01}+k_{10}) + k_{01}k_{10}(1-z)}.$$

We can thus find $G(z,t)$ by taking the inverse Laplace transform of $\tilde{G}(z,s)$ so that:

$$\forall z \text{ s.t. } |z| < 1, \quad G(z,t) = \mathcal{L}^{-1}\left\{ \frac{s + k_{10} + k_{01}}{s^2 + s(k_{01}+k_{10}) + k_{01}k_{10}(1-z)} \right\}.$$

Now, let us express by

$$s_{\pm}(z) = \frac{-(k_{10}+k_{01}) \pm \sqrt{(k_{10}+k_{01})^2 - 4\,k_{10}k_{01}(1-z)}}{2},$$

the two roots of the second degree equation in the denominator above. By now setting $\varepsilon := k_{01}/k_{10} \ll 1$ and $\delta := -2\epsilon + 4\epsilon z + \epsilon^2$ we have that:

$$k_{10}\sqrt{(1+\epsilon)^2 - 4\epsilon(1-z)} = k_{10}\sqrt{1+\delta} = k_{10}\left(1 - \epsilon + 2\epsilon z + O(\epsilon^2)\right).$$

whence we find:

$$s_+(z) = -\epsilon(1-z)\,k_{10} + O(\epsilon^2 k_{10}) = -k_{01}(1-z) + O(\epsilon\, k_{01}), \quad s_-(z) = -\big[1 + O(\epsilon)\big]\,k_{10}.$$

**Neglecting the fast-mode contributions.** Writing now

$$\widetilde{G}(z,s) = \frac{s + k_{10} + k_{01}}{(s - s_+(z))(s - s_-(z))} = \frac{C}{s - s_+(z)} + \frac{D}{s - s_-(z)},$$

we have that by direct computations we can compute

$$C = 1 + O(\epsilon), \qquad D = O(\epsilon).$$

In the time domain, we thus have

$$G(z,t) = C\,e^{s_+(z)t} + D\,e^{s_-(z)t} = e^{-k_{01}(1-z)t}\big[1 + O(\epsilon)\big] + O(\epsilon)\,e^{-k_{10}t}.$$

Therefore, for any $t$ such satisfying $k_{10}t \gg 1$, we have that $e^{-k_{10}t}$ is exponentially small, hence we may drop the second term in the expression above, thus getting:

$$G(z,t) \approx \exp\big[-k_{01}(1-z)t\big] = \sum_{n=0}^{\infty} \frac{(k_{01}t)^n}{n!} z^n,$$

i.e., for $t \gg 1/k_{10}$ the following approximation holds $N(t) \sim \mathrm{Poisson}(k_{01}t)$.

In fluorescence settings where $k_{10} \approx 10^9\,\mathrm{s}^{-1}$, the Poisson approximation becomes valid for any observation time more than a nanosecond. Beyond this timescale, the contribution of the fast relaxation mode $e^{-k_{10}t}$ becomes negligible, and the excitation–relaxation cycles effectively behave as independent, memoryless events. As a result, the photon count statistics converge rapidly to a Poisson distribution with rate $k_{01}$, and deviations from this model are only significant at extremely short timescales.

### 3 Supplementary Note 3: Optical and Camera System Modeling

The image formation process in fluorescence microscopy involves optical blur, spatial sampling, and detector-specific noise corruption. MUFASA simulates these effects through a modular pipeline, modeling both the physical optics, with a simplified model, and the camera electronics.

**Optical blur and spatial undersampling.** Photon emission at high spatial resolution is first blurred by the optical point spread function (PSF), modeled here as a 2D Gaussian $\mathcal{G}(\sigma)$, where $\sigma$ is a user-defined parameter that defines the spread of the PSF in pixel units. Let $\mathbf{x}^t \in \mathbb{R}^{H\times W}$ denote the high-resolution ground-truth image at time $t$. The blurred image is given by:

$$\mathbf{X}^t_{\text{blur}} = \mathcal{G}(\sigma) * \mathbf{x}^t,$$

where $\mathbf{X}^t_{\text{blur}} \in \mathbb{R}^{H\times W}$ is the 2D blurred image. If needed for downstream processing, this result is then flattened as a vector:

$$\mathbf{x}^t_{\text{blur}} = \Psi\mathbf{x}^t = \text{vec}(\mathbf{X}^t_{\text{blur}}).$$

To reflect the finite pixel size of the detector, the high-resolution image is spatially undersampled by a factor $L$, corresponding to a pixel area of $L \times L$ in the high-resolution grid. This operation is represented by a linear downsampling operator $U \in \mathbb{R}^{m\times n}$, where $n = H \times W$ is the number of high-resolution pixels and $m = n/L^2$ is the number of detector pixels. Each low-resolution pixel value is computed as the sum of intensities within its corresponding $L \times L$ region of the high-resolution image.

In the context of time-varying molecular emission, the final recorded intensity in each detector pixel integrates the blurred photon contributions from all fluorophores that were active during the acquisition time and spatially located within the corresponding region. This yields the low-resolution intensity:

$$\tilde{\mathbf{x}}^t = U\Psi\mathbf{x}^t + \mathbf{b},$$

where $\mathbf{b}$ models background fluorescence coming from out-of-focus regions.

**Photon detection and camera modeling.** ot all emitted photons reach or are registered by the detector due to optical losses, quantum efficiency limitations, and electronic artifacts. This induces the stochasticity of photon detection given by a Poisson distribution. The number of detected photoelectrons $n_{\text{ie}}$ is, then, modeled as:

$$n_{\text{ie}} = \mathcal{P}(QE \cdot \tilde{\mathbf{x}}^t + c),$$

where $QE$ is the quantum efficiency and $c$ is the clock-induced charge, a small spurious signal generated during charge transfer in the detector.

.

MUFASA supports multiple camera types:

- **CCD:** [6] Direct conversion with Gaussian readout noise:

$$n_{\text{oe}} = n_{\text{ie}} + \mathcal{N}(\mu, \sigma^2).$$

- **EMCCD:** [6] Stochastic amplification with a Gamma distribution, followed by Gaussian noise:

$$n_{\text{oe}} = \Gamma(n_{\text{ie}}, EM_{\text{gain}}) + \mathcal{N}(\mu, \sigma^2).$$

- **CMOS:** [11] Nonlinear pixel-specific gain function $g_p(\cdot)$, followed by Gaussian readout noise:

$$n_{\mathrm{oe}} = g_p(n_{\mathrm{ie}}) + \mathcal{N}(\mu, \sigma^2).$$

**Digitization.** The final output is quantized into Analog-to-Digital Units (ADUs):

$$\mathbf{y}^t = \min\left( \left\lfloor \frac{n_{\mathrm{oe}}}{e_{\mathrm{ADU}}} \right\rfloor + \mathrm{BL}, \mathrm{ADC}_{\mathrm{max}} \right),$$

where:

- $e_{\mathrm{ADU}}$ is the electron-to-ADU conversion factor,
- BL is the baseline offset,
- $\mathrm{ADC}_{\mathrm{max}}$ is the digital saturation level (e.g., 65535 for a 16-bit ADC).

This complete modeling chain ensures that simulated data replicates actual real fluorescence microscopy acquisitions.

**Denoising Acquisition Signal** To provide a rough estimation of the underlying photon emission signal $\tilde{\mathbf{x}}^t$ from the observed digitized acquisition $\mathbf{y}^t$, we exploit the known structure of the physical acquisition chain detailed previously. This knowledge allows us to leverage the statistical properties of each noise source—additive Gaussian readout noise, gamma-distributed amplification noise (for EMCCDs), and Poisson-distributed photon counting noise—to denoise the signal in a modular fashion. Each noise component is addressed with methods tailored to its specific distribution.

All camera types include a Gaussian noise component at the readout stage, modeled as:

$$n_{\mathrm{oe}} = \hat{n} + \mathcal{N}(0, \sigma^2),$$

where $\hat{n}$ is the intermediate signal prior to quantification. To remove such noise, we use Gaussian denoisers such as wavelet-based methods to estimate $\hat{n}$ and suppress readout noise, yielding a denoised signal $\hat{n}^{(1)}$.

For EMCCD sensors, after Gaussian denoising, we normalize by the known EM gain to approximate the input to the amplifier:

$$\hat{n}_{\mathrm{ie}} \approx \frac{\hat{n}^{(1)}}{\mathrm{EM}_{\mathrm{gain}}}.$$

For CMOS cameras, note that a nonlinear, pixel-specific gain function $g_p(\cdot)$ is applied. The calibration data is generally available for CMOS, so we invert this transformation numerically:

$$\hat{n}_{\mathrm{ie}} \approx g_p^{-1}(\hat{n}^{(1)}).$$

In all cases, the final signal $\hat{n}_{\mathrm{ie}}$ is modeled as a realization of Poisson-distributed photoelectrons:

$$n_{\mathrm{ie}} \sim \mathcal{P}(QE \cdot \tilde{x}^t + c),$$

for which a Poisson denoiser $\mathcal{D}_{\mathrm{Pois}}$, such as PURE-LET [7] can be applied, to estimate a smoother signal:

$$\hat{n}_{\mathrm{denoised}} = \mathcal{D}_{\mathrm{Pois}}(\hat{n}_{\mathrm{ie}}).$$

We retain $\hat{n}_{\text{clean}}$ as a denoised representation of the observed photon counts.

This physics-aware, layered denoising pipeline provides a principled way to stabilize and enhance noisy acquisitions, and is used as a preprocessing step throughout MUFASA-Design.

### 4 Supplementary Note 4: SOFI-Simlulation Tools

SOFI Simulation Tools [5] uses a continuous-time Markov model with a state space $|\mathcal{S}| = 2$ with two regimes—"on" and "off"—and models transitions between them using exponential holding times. The rates $k_{\text{on}}$ and $k_{\text{off}}$ are defined by the mean durations in each state:

$$k_{\text{on}} = 1/T_{\text{on}}, \quad k_{\text{off}} = 1/T_{\text{off}}$$

. During the "on" regime, SOFI assumes a fixed photon emission rate $I_{\text{on}}$, so that the total number of photons emitted during a "on" event is:

$$N_{\text{photon}}^{\text{SOFI}} = I_{\text{on}} \cdot t_{\text{on}}, \quad \text{with} \quad t_{\text{on}} \sim \mathcal{E}(T_{\text{on}})$$

This stochastic framework allows simplified simulations of fluorophore blinking, which is valuable for generating synthetic image stacks for testing super-resolution optical fluctuation imaging (SOFI) and balanced SOFI (bSOFI) pipelines. The simulator also includes tunable parameters for choosing fluorophore density, PSF shape, bleaching dynamics, noise statistics (e.g., Poisson photon shot noise, Gaussian camera readout), and supports both SOFI and STORM reconstruction algorithms. Importantly, the bSOFI module applies a linearization step to correct the nonlinear brightness dependence inherent in higher-order cumulants. For further technical details, the full simulation architecture is described in [5].

### 5 Supplementary Note 5: Camera Configurations

The configurations listed in Table 1 are representative examples. Our framework is designed to be modular,allowing any protocol to be simulated with any of the camera models to study the impact of different hardware choices. These configurations correspond to the frames examples given in the main results and the supplementary notes.

**Supplementary Table 1. Camera configurations used for each simulated SRM protocol in MUFASA-Sim.**

| **Parameter** | **PALM** | **STORM** | **FF** | **Blinking** |
|---|---|---|---|---|
| Camera type | CCD | CCD | EMCCD | CMOS |
| Quantum efficiency (QE) | 0.9 | 0.9 | 0.9 | 0.9 |
| Read noise $\sigma_R$ ($e^-$) | 70.58 | 70.58 | 60.58 | 1.58 |
| Clock-induced charge $c$ ($e^-$) | 0.002 | 0.002 | 0.002 | 0.002 |
| Electron/ADU $e_{\text{ADU}}$ | 10.04 | 100.04 | 0.74 | 20.74 |
| Baseline (BL) (ADU) | 367 | 367 | 367 | 367 |
| ADC max | 65535 | 65535 | 65535 | 65535 |
| EM (Amplification) | - | - | 500 | - |
| PSF FWHM (nm) | 1200 | 1200 | 308 | 308 |
| Pixel size (nm) | 100 | 100 | 100 | 100 |
| Background (photons) | 20 | 20 | 20 | 20 |
| CMOS gain function $g_p(e)$ | – | – | – | $\begin{cases} 0.9e, & e < 200 \\ 180 + 0.4(e - 200), & 200 \leq e < 1000 \\ 340 + 0.1(e - 1000), & e \geq 1000 \end{cases}$ |

### 6 Supplementary Note 6: Experimental Protocol Parameters

**Supplementary Table 2. Experimental setup parameters for each protocol.**

| Parameter | PALM | STORM | FF | Blinking |
|---|---|---|---|---|
| Experiment Duration (s) | 1.00 | 1.00 | 1.00 | 1.00 |
| Number of Frames | 500 | 500 | 100 | 100 |
| Frame Length (ms) | 10 | 10 | 10 | 10 |
| Excitation Power (W/cm$^2$) | 5 | 7 | 2 | 3 |
| Activation Rate (s$^{-1}$) | 1 | - | – | – |
| Excitation Wavelength (nm) | 647 | 647 | 647 | 647 |

**Supplementary Table 3. Molecular parameters used for each protocol.**

| Parameter | PALM | STORM | FF | Blinking |
|---|---|---|---|---|
| Extinction Coefficient ($\epsilon$) (M$^{-1}$cm$^{-1}$) | 270000 | 270000 | 270000 | 270000 |
| Excitation Lifetime ($\tau$) (s) | $2. \cdot 10^{-9}$ | $2 \cdot 10^{-9}$ | $2 \cdot 10^{-9}$ | $2 \cdot 10^{-9}$ |
| Cycles Before Bleaching | $5 \cdot 10^5$ | $5 \cdot 10^5$ | $5 \cdot 10^3$ | $1 \cdot 10^3$ |
| Non-Radiative Decay Rate ($\alpha_{nr}$) | 10 | 2 | 1 | 3 |
| ISC Rate ($\alpha_{isc}$) | $1 \cdot 10^{-2}$ | $5 \cdot 10^{-4}$ | $1 \cdot 10^{-5}$ | $5 \cdot 10^{-4}$ |

### 7 Supplementary Note 7: Extracting Single Molecule Signals from Experimental Acquisitions

To evaluate the accuracy of MUFASA on real data, we considered time-lapse fluorescence microscopy acquisitions of a sample labeled with EGFP (Enhanced Green Fluorescent Protein). The properties of EGFP are summarized in Table 4.

**Supplementary Table 4. Photophysical properties of EGFP used for experimental validation.**

| | |
|---|---|
| Excitation Wavelength ($\lambda_{\text{exc}}$) | 488 nm |
| Emission Wavelength ($\lambda_{\text{em}}$) | 507 nm |
| Extinction Coefficient ($\epsilon$) | 55,900 $\text{M}^{-1}\text{cm}^{-1}$ |
| Quantum Yield (QY) | 0.6 |
| Fluorescence Lifetime | 2.6 ns |

The experiment consists of recording a 512 × 512 pixel time series of 250 frames using an EMCCD camera. Each frame corresponds to an amplified digital signal. Prior to illumination, a background frame was acquired to estimate the camera intrinsic noise characteristics (Figure 1, left). Subsequent frames captured the fluorescent dynamics of EGFP-tagged molecules under continuous illumination (Figure 1, right).

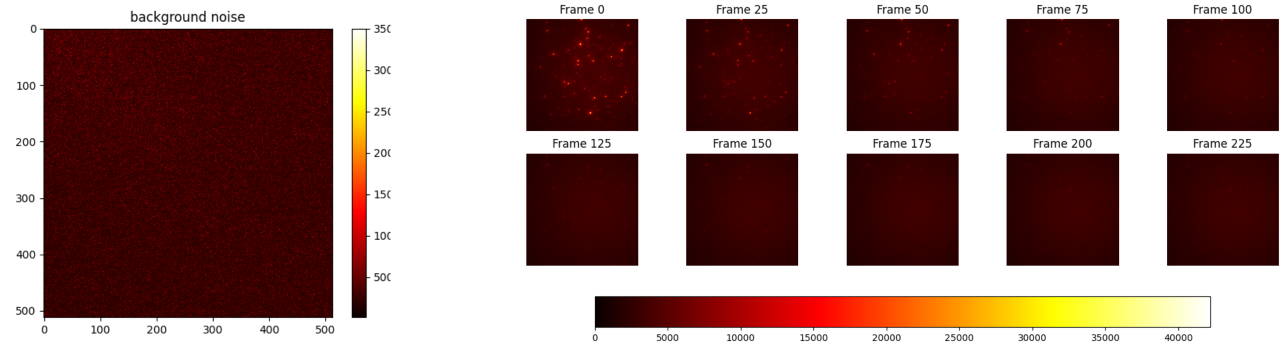


**Supplementary Figure 1. Camera output before and during fluorescence acquisition. Left:** Background noise frame recorded before illumination. **Right:** Fluorescence signal after illumination of EGFP-expressing sample at different time frames. (every 2.5s).

We applied a threshold to each frame to isolate pixels with intensities significantly above the background noise, identifying candidate locations likely to contain active fluorophores. For each candidate region, we assumed that the brightest pixel corresponds to the molecule's spatial center, with the surrounding intensity arising from optical blur (PSF spread). We then extracted the full temporal intensity profile from this central pixel across all frames. Signals that did not exhibit the expected fluorescence dynamics were discarded, ensuring that only well-isolated single-molecule traces were retained.

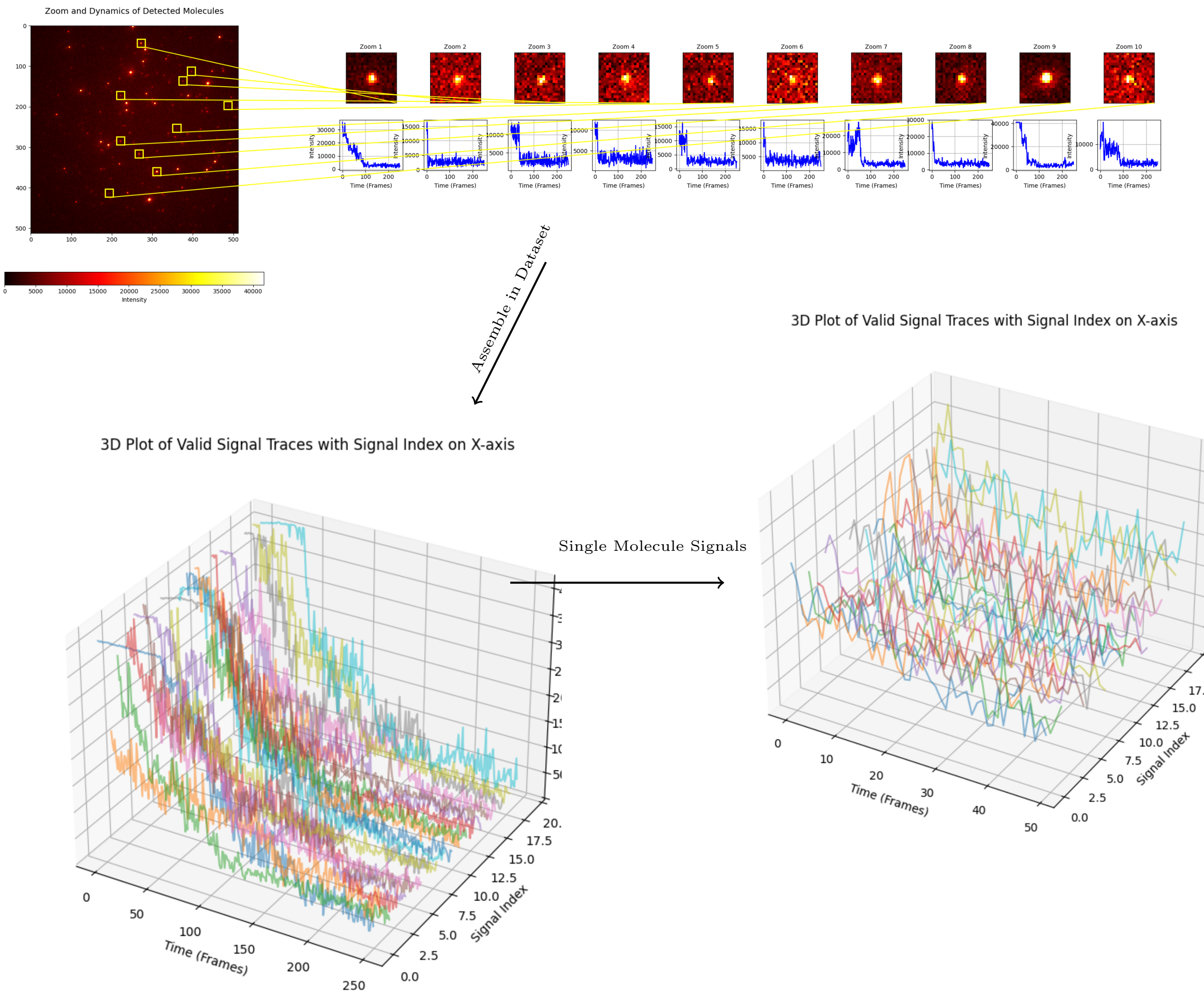


**Supplementary Figure 2. From full-frame fluorescence to single-molecule signal extraction. Top:** Raw intensity dynamics from all candidate pixels exceeding the background noise threshold. For each region, the brightest pixel is assumed to correspond to the fluorophore center, and its temporal trace is extracted. **Bottom left:** Segmented traces of candidate molecules across the entire acquisition sequence (250 frames). **Bottom right:** Final cleaned dataset of individual single-molecule fluorescence signals, obtained by selecting a time window of 50 frames just before bleaching, during which all chosen emitters remain active. This ensures a consistent set of unbleached, spatially isolated single-molecule trajectories suitable for model validation.

This procedure enabled the extraction of a dataset consisting of 20 single-molecule fluorescence traces across 50 frames each, suitable for validation of simulation pipelines.

### 8 Supplementary Note 8: Experimental Conditions and Parameter Inference for STORM Validation

To reproduce the experimental conditions for STORM validation, we used data from the Zenodo repository "STORM Vectashield datasets (Tubulin)" [8]. Specifically, we extracted acquisition metadata and sample preparation details from the associated dataset description, which included the following:

- **Excitation wavelength:** 641 nm laser (Coherent, CUBE 640–100C)
- **Excitation intensity on sample:** ∼1–2 kW/cm$^2$
- **Camera:** Andor iXonEM+ (EMCCD) operated in conventional CCD mode
- **Pixel size:** Effective 100 nm
- **Objective:** Olympus 100x 1.3NA, with 1.6x magnification lens

All simulations used fluorophores modeled as Alexa Fluor 647 with known spectral and photophysical properties from datasheets. While excitation rate $k_{01}$ was derived from the given parameters, some transition rates were not directly available, hence they had to be estimated:

- **Intersystem crossing rate $k_{1T}$:** Tuned within a plausible range (based on $\phi_{ISC} \in [0.001, 0.01]$) to match the characteristic dark-state durations observed experimentally.
- **Triplet state decay rate $k_{T0}$:** Adjusted to yield temporal sparsity consistent with recorded emission bursts.
- **Photobleaching rate $k_{1B}$:** Calibrated to reproduce the observed signal decay across 5000 frames.

We stress that the goal was not to fit data exactly, but to match the overall statistics and event dynamics under reported environmental and instrumental conditions. The camera model (including EMCCD noise) used the manufacturer's spec sheet `Spec-Sheet_iXon.JPG`, as detailed in Supplementary Note 3.

All resulting simulation parameters are available in Supplementary Table 5.

| **Parameter** | **Value** | **Description / Source** |
|---|---|---|
| | **Experimental Setup** | |
| Microscopy protocol | STORM | Single-molecule localization microscopy |
| Excitation wavelength $\lambda$ | 641 nm | Laser from Zenodo metadata [8] |
| Laser power density $P$ | 1,000 W/cm² | Conservative estimate (range: 1–2 kW/cm²) |
| Frame duration | 36 ms | Exposure time per frame |
| Number of frames | 2,000 | Full acquisition duration |
| Camera | Andor iXonEM+ | Effective pixel size: 100 nm |
| | **Fluorophore: Alexa Fluor 647** | |
| Molar extinction coefficient $\varepsilon$ | 239,000 $M^{-1}cm^{-1}$ | Thermo Fisher datasheet |
| Excited-state lifetime $\tau_1$ | 1 ns | Typical for Alexa 647 |
| Intersystem crossing coefficient $\alpha_{ISC}$ | $1.6 \times 10^{-3}$ | Calibrated for matching OFF durations |
| Triplet decay rate $\alpha_{nr}$ | 0.1 $s^{-1}$ | Tuned to reproduce dark-state returns |
| Cycles before bleaching | $10^9$ | Effective; negligible bleaching during sim |

**Supplementary Table 5.** Experimental and photophysical parameters used for MUFASA-STORM simulation.

### 9 Supplementary Note 9: Wasserstein Distance $W_1$

To quantitatively assess the performance of the MUFASA simulations, we consider a validation framework based on 1-Wasserstein distance ($W_1$) [12]. Given the interpretation of our measurements as realisations of stochastic processes, we use this quantity as a measure of discrepancy between the distributions of photon emissions from simulated and experimentally observed datasets.

Specifically, we compute $W_1$ on concatenated photon emission traces from multiple fluorophores. Namely, given $N$ molecules, the emission time series from each molecule (both simulated and experimental) are concatenated into two large arrays:

$$\mathbf{X}_{\text{sim}} = [x_1^{\text{sim}}, x_2^{\text{sim}}, \dots, x_N^{\text{sim}}] \quad \text{and} \quad \mathbf{X}_{\text{exp}} = [x_1^{\text{exp}}, x_2^{\text{exp}}, \dots, x_N^{\text{exp}}],$$

where, for $i = 1, \dots, N$ each $x_i^{\text{sim}} \in \mathbb{R}^T$ represents the photon count time series for the $i^{\text{th}}$ fluorophore over $T$ timepoints. The independence of emission events ensures that concatenating photon emission traces is a valid computational procedure capturing the overall statistical behavior across multiple fluorophores.

More precisely, by considering the empirical measures

$$\mu = \frac{1}{M} \sum_{i=1}^{M} \delta_{x_i^{\text{sim}}}, \qquad \nu = \frac{1}{M} \sum_{j=1}^{M} \delta_{x_j^{\text{exp}}},$$

both having uniform weights, the computation of the 1–Wasserstein distance reduces to consider the optimal-transport linear program:

$$W_1(\mu, \nu) = \min_{\Gamma \in \mathbb{R}_+^{M \times M}} \sum_{i,j=1}^{M} \Gamma_{ij} \, C_{ij}, \qquad \sum_j \Gamma_{ij} = \tfrac{1}{M}, \; \sum_i \Gamma_{ij} = \tfrac{1}{M},$$

where the **cost matrix** has entries

$$C_{ij} = \|x_i^{\text{sim}} - x_j^{\text{exp}}\|.$$

A lower $W_1$ indicates closer alignment between simulated and experimental emission distributions, thereby validating the accuracy and realism of the simulation framework.

**Practical computation** To compute this distance efficiently, we use the Python Optimal Transport (POT) library [4], which implements a range of optimal transport solvers. Specifically, we employ the exact linear programming solver *ot.emd2*:

```python
import ot
wass_dist = ot.emd2(a, b, C)
```

- `a` and `b` are uniform histograms over the rows of `X_sim` and `X_exp` respectively.
- `C` is the cost matrix defined above.

### 10 Supplementary Note 10: Quantitative assessments via Wasserstein-distance of FF dynamics

We evaluated MUFASA's ability to replicate single-molecule emission statistics under the FF protocol by comparing its simulated photon count distributions to experimentally measured data. The experimental dataset was acquired using an EMCCD camera imaging EGFP-labeled samples under continuous illumination, yielding amplified digital signals representative of photon emission time series. Details of molecule preparation, acquisition protocols, and the single-molecule signal extraction pipeline are provided in [10] and in Supplementary Note 7 .

To quantitatively assess the simulator's ability to reproduce real-world fluorescence dynamics, we computed the first-order Wasserstein distance, as described in Supplementary Note 9, between the distribution of experimentally observed photon counts per frame and the outputs of three simulation strategies: MUFASA, a standard Poisson process model, and SOFI Tools.

The Poisson process model was included as a baseline, as it represents the simplest counting process: photon arrivals modeled as a homogeneous Poisson process with a constant rate $\mu$, derived from theoretical considerations. Specifically, we used the closed-form excitation rate $k_{01}$ defined in Equation ??, and set $\mu = k_{01} \cdot \Delta t$, with $\Delta t$ being the duration of a frame. This model assumes memoryless photon emission with no sub-state variability, consistent with common assumptions in FF-SRM theory.

Parameter selection for all simulations, including MUFASA and SOFI Tools, was based on the experimental acquisition conditions described earlier. For the FF protocol, no additional parameter tuning was required beyond the known values of illumination power, wavelength, fluorophore properties (e.g., $\epsilon$), and quantum yield. These inputs were sufficient to determine the necessary transition rates for MUFASA via closed-form expressions. For the SOFI Tools baseline, which assumes a two-state Markov model with constant photon emission during the "on" state, we used values extracted from MUFASA simulations to ensure alignment. Specifically, one of its parameters is the emission intensity, which was set equal to the maximum value observed in the MUFASA-generated photon trace. This approach ensures a fair comparison, aligning both simulators with respect to photophysical timescales and emission amplitudes.

The Wasserstein distance provides a meaningful metric for quantifying the difference between two probability distributions, capturing both the distributional shape and the geometry of the data. In our context, we are comparing stochastic photon emission traces. Metrics such as Euclidean distance (e.g., $\ell_2$ norm) are not well suited for this task, as they treat the signals deterministically and penalize pointwise deviations without accounting for underlying statistical variability. Kullback–Leibler (KL) divergence can be undefined or infinite when the two distributions have non-overlapping supports, making it unstable for comparing empirical or noisy data [2]. Photon emission time traces from different simulators may yield distributions where one has a tail or concentration that the other doesn't, especially with noise. KL may diverge or be unstable, but Wasserstein gives a robust, interpretable difference in such cases.

Although MUFASA achieves strong agreement with the experimental data, its Wasserstein distance is slightly elevated. This can be attributed to the explicit modeling of EMCCD cameras, where the stochastic amplification of photoelectrons introduces additional variability. Notably, one can analytically demonstrate that the Wasserstein distance scales linearly with the amplification factor in EMCCD systems.

As shown in Supplementary Fig. 3, MUFASA achieved a Wasserstein distance of $W_1 = 2263$, closely aligning with the experimental data and marginally outperforming the Poisson model ($W_1 = 2269$). By contrast, SOFI Tools produced a substantially higher distance ($W_1 = 3065$), indicating a poorer fit to the empirical distribution. This deviation reflects the limitations of assuming constant photon emission within on-periods, which fails to capture the intrinsic variability modeled by MUFASA. These results underscore MUFASA's capacity to simulate realistic photophysical behavior, capturing both the stochastic switching and emission-level fluctuations observed in experimental FF data.

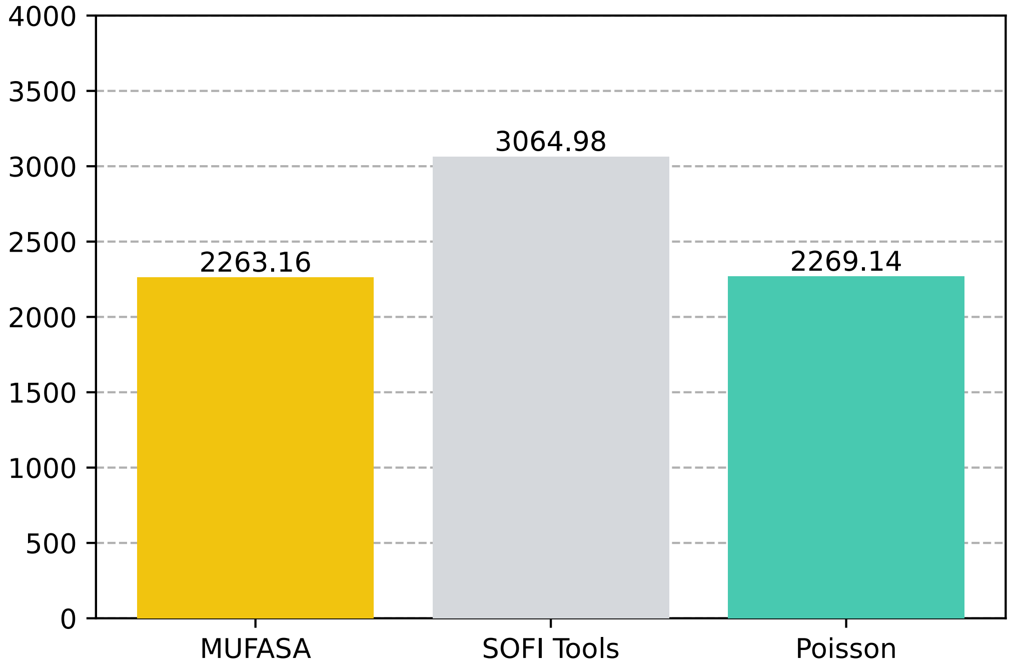


**Supplementary Figure 3. Comparison of FF simulations using the 1-Wasserstein distance.** Wasserstein distances ($W_1$) between experimental photon count distributions and three simulation models: MUFASA, Poisson, and SOFI Tools. Lower values indicate closer alignment with the empirical distribution of experimental data. MUFASA achieves the best match, highlighting its capacity to capture realistic emission variability under the FF protocol.

### 11 Supplementary Figures

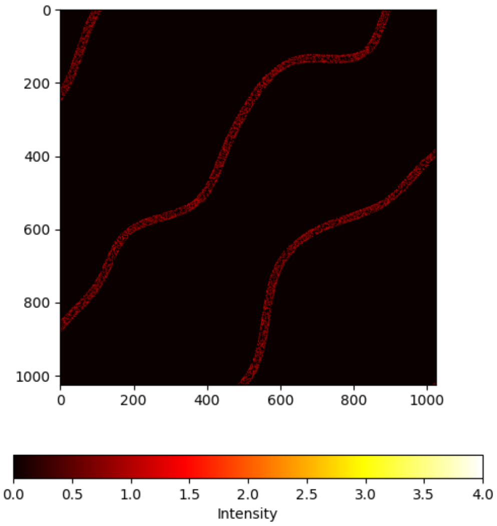


**Supplementary Figure 4. Fluorophore grid layout.** Arrangement of 6,498 fluorophores on a $2048 \times 2048$ grid used for input simulation.

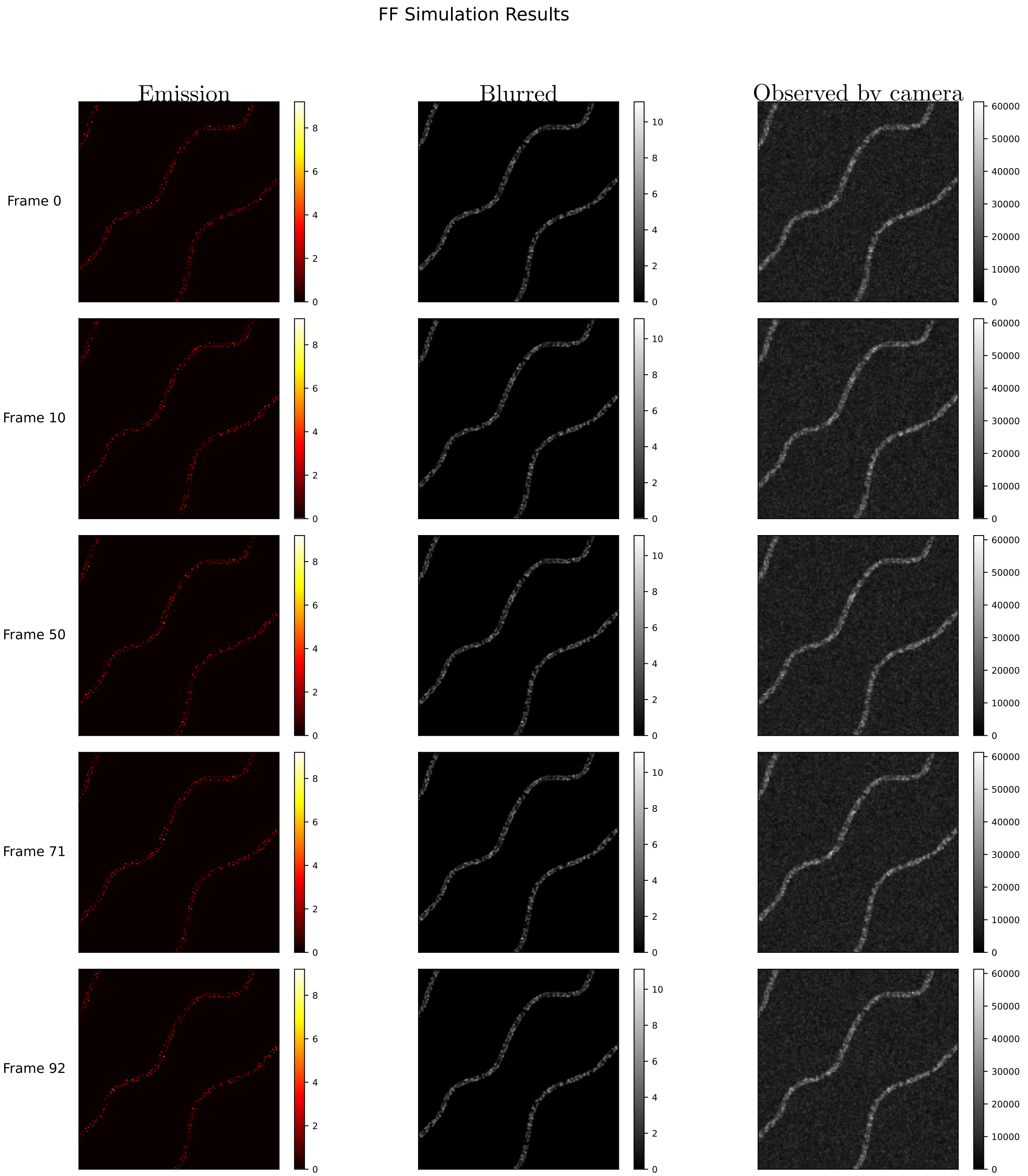


**Supplementary Figure 5. MUFASA-Sim FF protocol output.** Selected frames (0, 10, 50, 71, 92) from a 100-frame fluorescence simulation. **Left column**: "Emission" maps showing per-pixel photon emission rate in photons per frame (0–6 photons/frame color-scale). **Middle column**: "Blurred" images after applying the microscope point-spread function , in intensity units. **Right column**: "Observed by camera" output including shot and read noise, in camera digital units (ADU; 0–60,000 ADU scale). The sequence demonstrates continuous and spatially consistent photon emission and detection throughout the simulation protocol.

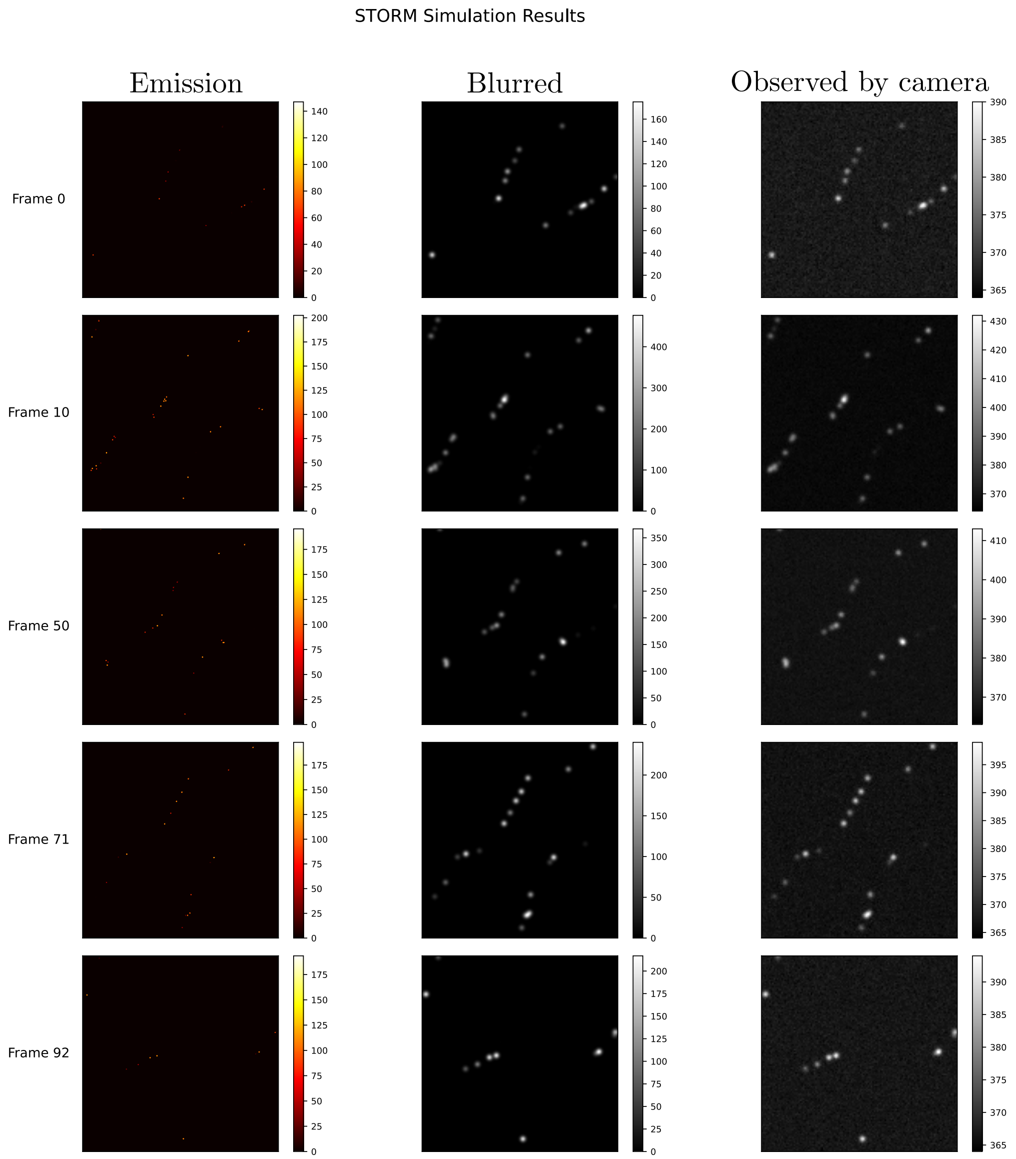


**Supplementary Figure 6. MUFASA-Sim STORM protocol output.** Selected frames (0, 10, 50, 71, 92) from a 500-frame stochastic blinking simulation. **Left column**: "Emission" maps showing stochastic photon emission events in photons per frame (0–200 photons/frame color-scale). **Middle column**: "Blurred" images after applying the point-spread function, in intensity units (0–200). **Right column**: "Observed by camera" output including shot and read noise, in analog-to-digital units (ADU; 0–400 ADU scale). The output reflects characteristic blinking behavior typical of STORM imaging.

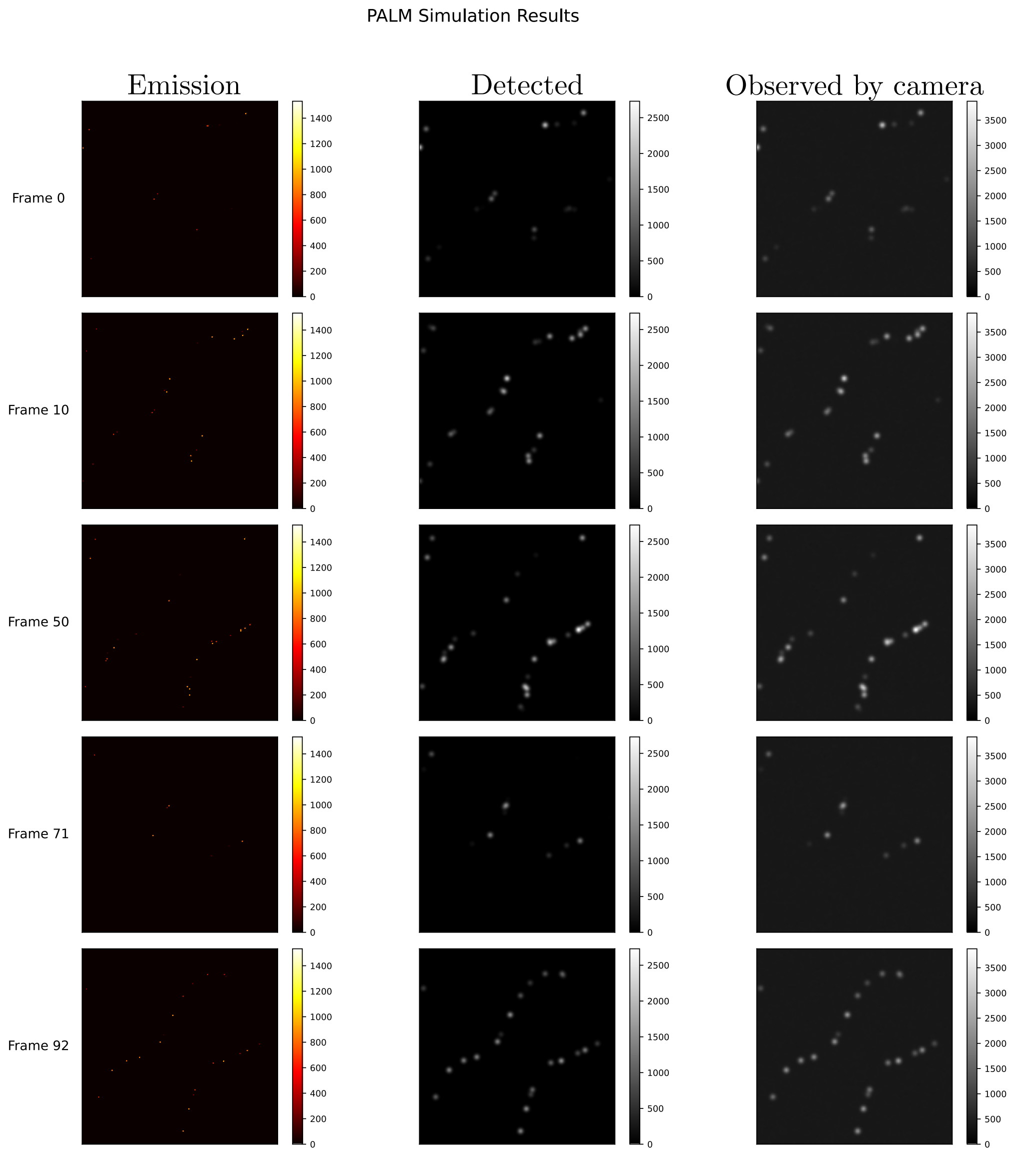


**Supplementary Figure 7. MUFASA-Sim PALM protocol output.** Selected frames (0, 10, 50, 71, 92) from a 500-frame simulation of photoactivated localization microscopy. **Left column**: "Emission" maps showing photon bursts from activated fluorophores in photons per frame (0–1400 photons/frame color-scale). **Middle column**: "Detected" signals after applying the point-spread function, in intensity units (0–2500). **Right column**: "Observed by camera" output including noise and digitization, in analog-to-digital units (ADU; 0–2600 ADU scale). The simulation highlights sparse, stochastic activation consistent with PALM imaging.

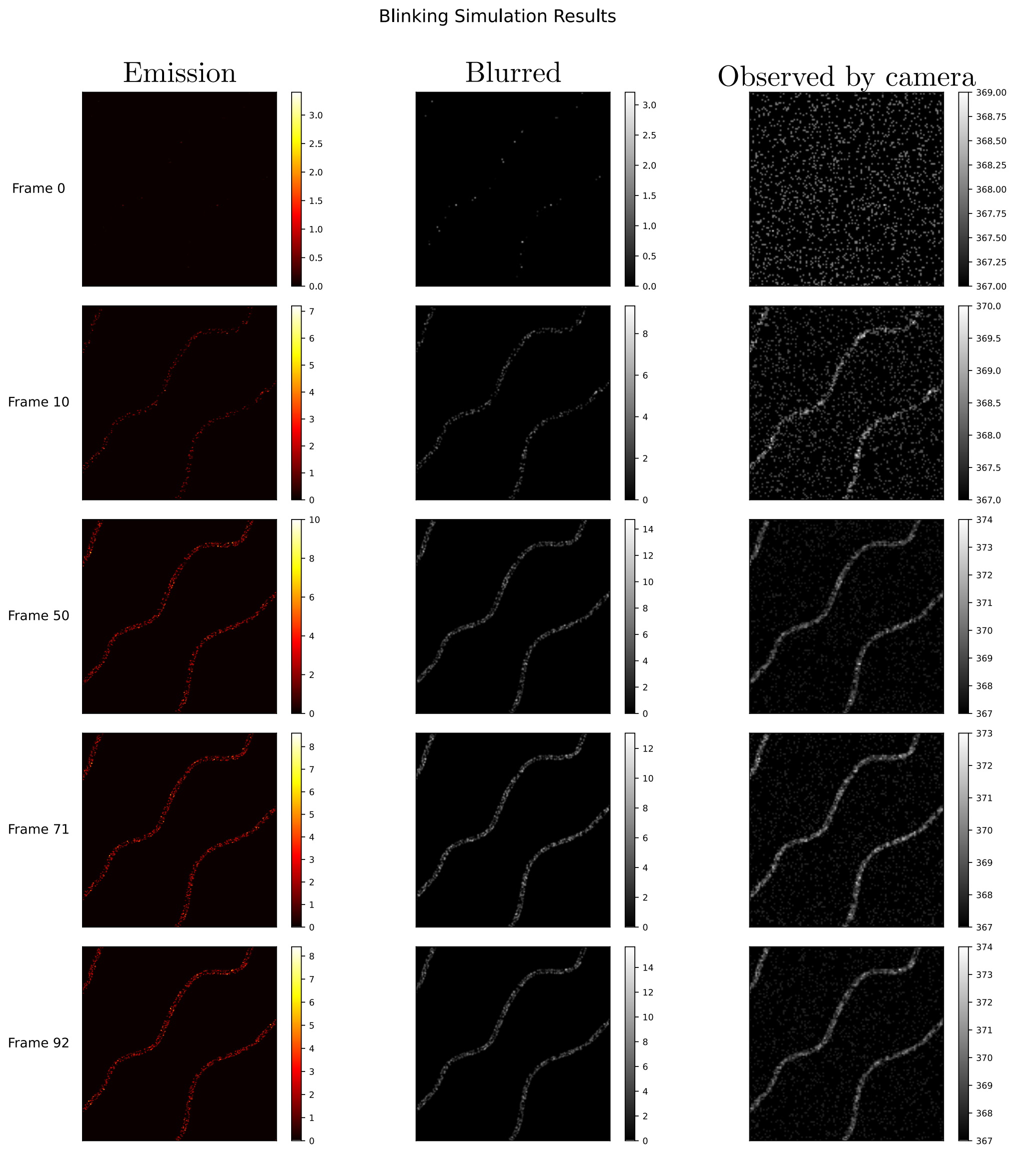


**Supplementary Figure 8. MUFASA-Sim Blinking protocol output.** Selected frames (0, 10, 50, 71, 92) from a 100-frame blinking simulation. **Left column**: "Emission" maps showing frame-wise variation in emission in photons per frame (0–10 photons/frame color-scale). **Middle column**: "Blurred" images after applying the point-spread function, in normalized intensity units (0–14). **Right column**: "Observed by camera" output including camera noise, in analog-to-digital units (ADU; 0–380 ADU scale). The sequence demonstrates stochastic activation and deactivation of emitters across time.

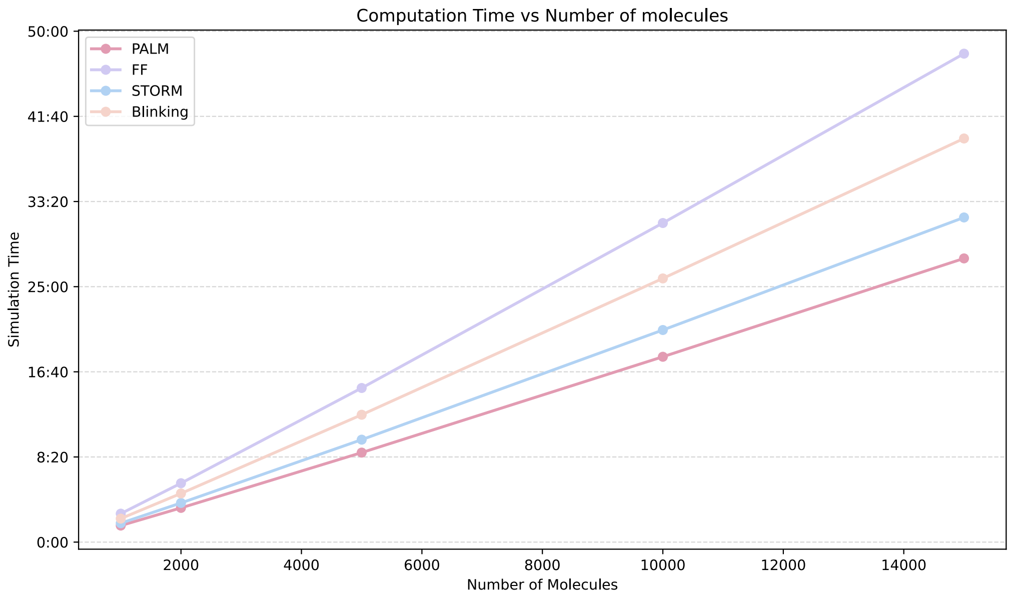


**Supplementary Figure 9. MUFASA Simulation runtime as a function of the number of fluorophores across four imaging protocols.** Computational runtimes for simulating PALM, STORM, FF, and Blinking protocols increase approximately linearly with the number of fluorophores. Times were obtained from actual simulations performed with 1,000, 2,000, 5,000, 10,000, and 15,000 molecules for each protocol under identical conditions. PALM and STORM exhibited shorter runtimes due to the early bleaching of the molecules, while FF and Blinking incurred longer runtimes.

### References


[1] Robert Alicki and Karl Lendi. *Quantum Dynamical Semigroups and Applications*. Lecture Notes in Physics. Springer Berlin, Heidelberg, 1 edition, April 2007. Hardcover ISBN: 978-3-540-70860-5, Published: 23 April 2007. Softcover ISBN: 978-3-642-08985-5.

[2] Martin Arjovsky, Soumith Chintala, and Léon Bottou. Wasserstein generative adversarial networks. In Doina Precup and Yee Whye Teh, editors, *Proceedings of the 34th International Conference on Machine Learning*, volume 70 of *Proceedings of Machine Learning Research*, pages 214–223. PMLR, 06–11 Aug 2017.

[3] Jurek W. Dobrucki and Ulrich Kubitscheck. *Fluorescence Microscopy*, chapter 3, pages 85–132. John Wiley & Sons, Ltd, 2017.

[4] Rémi Flamary, Nicolas Courty, Alexandre Gramfort, and et al. Pot: Python optimal transport. *Journal of Machine Learning Research*, 22(78):1–8, 2021.

[5] Arik Girsault, Tomas Lukes, Azat Sharipov, Stefan Geissbuehler, Marcel Leutenegger, Wim Vandenberg, Peter Dedecker, Johan Hofkens, and Theo Lasser. SOFI Simulation Tool: A Software Package for Simulating and Testing Super-Resolution Optical Fluctuation Imaging. *PLoS ONE*, 11(9):e0161602, 2016. PMCID: PMC5008722.

[6] M Hirsch, R.J. Wareham, M.L. Martin-Fernandez, M.P. Hobson, and D.J. Rolfe. A stochastic model for electron multiplication charge-coupled devices–from theory to practice. *PLoS One*, 8(1):e53671, Jan 2013. Epub 2013 Jan 31.

[7] Florent Luisier, Thierry Blu, and Michael Unser. Image denoising in mixed poisson–gaussian noise. *IEEE Transactions on Image Processing*, 20(3):696–708, 2011.

[8] Nicolas Olivier and Debora Keller. STORM Vectashield datasets (Tubulin). `https://doi.org/10.5281/zenodo.7620025`, 2023. Zenodo dataset.

[9] Sheldon M. Ross. *Introduction to Probability Models*. Academic Press, 11th edition, 2014. Chapter 6, pp. 371–375.

[10] Patrice Royal, Aurélie Andres-Bilbe, Patricia Ávalos Prado, Camille Verkest, Benjamin Wdziekonski, Samuel Schaub, Anne Baron, Florian Lesage, Xavier Gasull, Joshua Levitz, and Guillaume Sandoz. Migraine-associated tresk mutations increase neuronal excitability through alternative translation initiation and inhibition of trek. *Neuron*, 101(2):232–245.e6, January 2019. Epub 2018 Dec 17.

[11] Hui Tian and Abbas El Gamal. *Noise Analysis in CMOS Image Sensors*. Dissertation, Stanford University, Stanford, CA, 2000. Ph.D. dissertation.

[12] Cédric Villani. *Optimal transport: old and new*, volume 338. Springer, 2008.
